# Distributed Cortical Network Dynamics of Binocular Convergent Eye Movements in Humans

**DOI:** 10.1101/2025.08.15.670412

**Authors:** Farzin Hajebrahimi, Suril Gohel, Michael W. Cole, Tara L. Alvarez

**Author notes:** Corresponding Authors: Farzin Hajebrahimi, PhD: New Jersey Institute of Technology, Newark, New Jersey, Tara L Alvarez, PhD: New Jersey Institute of Technology, Newark, New Jersey. Funding: This publication was made possible, in part, by the National Eye Institute of the National Institutes of Health, Department of Health and Human Services, Bethesda, Maryland NEI-R01EY023261 (Alvarez). Competing interests: None declared.

## Abstract

Neuroimaging studies in humans have localized brain functions to specific brain regions, but a recent shift toward distributed network-based models of brain function promises deeper insights into the network processes that generate brain functionality. Resting-state functional connectivity provides a rich mapping of the brain’s network architecture, linking with both underlying structure and task-evoked responses across the whole brain. In this study, we utilized a model based on propagation of task-evoked activations over resting-state functional connectivity networks to identify cortical contributions to localized functional brain activations associated with binocular convergent eye movements. Binocular vision is crucial for daily routine activities, with its impairment leading to significant challenges in daily life. The distributed network-level mechanisms of binocular convergent eye movements remain unknown. Results showed that mapping activity flow over brain connections accurately generated actual brain activations associated with convergent eye movements, which were distinct from those observed during control tasks. The visual and dorsal attention networks dominated the propagation of activations through resting-state connections during convergent eye movements. Submodel analyses further revealed that restricting activity flow to individual networks, such as the visual or dorsal attention systems alone, substantially reduced model accuracy, underscoring the necessity of distributed, whole-brain contributions. In conclusion, highly distributed network pathways are involved in convergent eye movements, with some pathways contributing much more than others, providing important implications for future clinical models of binocular dysfunction.

## Introduction

Understanding how the brain generates behavioral responses is a primary objective of neuroimaging studies ^1^. Conventionally, this effort has concentrated on identifying brain activations in specific brain regions corresponding to particular motor or cognitive functions across various experimental conditions. Through stimulus-evoked or task-evoked functional magnetic resonance imaging (fMRI), many studies have characterized localized neural responses across domains such as visual, language, and motor control ^2–5^. This approach has established functional specialization as a core organizational principle of the brain, highlighting the roles of individual regions in supporting specific tasks.

Over the past two decades, however, a shift toward network-based models has redefined this view by emphasizing the importance of interregional interactions ^6–8^. A pivotal development in this direction was the introduction of resting-state functional connectivity, which revealed that spontaneous activity during rest is structured and reproducible, reflecting stable patterns of coordination across distant brain regions ^9–11^. These intrinsic connectivity patterns have been shown to align closely with task-evoked activation networks, offering a powerful framework for understanding how the brain supports cognition through dynamic, distributed interactions ^12–14^. Building upon this foundation, network neuroscience has extended its scope to evaluate structural, functional, and effective connectivity during both resting and task conditions ^6,7,15,16^, reflecting a broader shift in neuroscience toward understanding the brain as a networked system.

Furthermore, a broader framework emphasizes that task-evoked brain responses are not solely driven by activity within isolated regions, but also shaped by interactions across distributed networks ^15,17–21^. These distributed processes have been studied through task-evoked functional connectivity ^22^, which captures statistical dependencies between distant regions during task performance. Although some methodological challenges have been proposed ^23,24^, studies adopting this perspective have shown that many cognitive functions are supported by interactions across large-scale functional networks that dynamically reconfigure during task performance.

In parallel, accumulating evidence has demonstrated a close correspondence between structural and functional connectivity, particularly in resting-state networks ^25–27^, as well as similarity between resting-state and task-state functional connectivity ^13,28–31^. Because structural connections enable information transmission between regions, the anatomical organization of the brain is thought to shape functional interactions ^32^. This relationship suggests that the architecture of white matter pathways provides a foundational framework for shaping the patterns of interregional communication observed during both rest and task states. From this perspective, structural connectivity plays a fundamental role in determining task-evoked responses. In support of this view, studies have shown that a region’s structural connectivity profile can predict the spatial pattern of its task-related activations ^33,34^.

Furthermore, studies have expanded this perspective from structural connectivity to functional connectivity, aiming to predict task-evoked activations. Recent research has shown that task-specific brain activity reflects not only localized responses but also widespread network engagement ^21,35,36^. This points to a dynamic interplay between local processing and distributed neural interactions. Building on this, generative modeling approaches have demonstrated that regional task-evoked activity can be estimated based on resting-state functional connectivity patterns ^35,36^. One such framework, known as activity flow mapping, models how task activations may arise from the propagation of activity across distributed functional pathways ^35,37^. This method assumes that local activity is shaped, at least in part, by interactions with distal regions, reflecting a fundamentally network-driven process underlying cognitive function.

Previous research applying this framework has demonstrated that distributed connectivity patterns significantly contribute to visual processes such as category recognition during memory and perception tasks ^38^. This supports the idea that localized activations associated with visual processes emerge through both region-specific computations and broader network interactions. Therefore, large-scale brain network connectivity is proposed to serve as a fundamental underlying mechanism for the emergence of localized activations ^39^. In other words, the way local brain regions are connected through large-scale networks is crucial in determining how these regions respond to a specific task.

This activity flow account may extend to more fundamental sensorimotor functions, including oculomotor control, where network-level mechanisms could play a similar role. Since activity flow models utilize empirical data as model parameters, this approach could provide a strong explanation of such functions and further test that explanation with observed empirical activations associated with the oculomotor control system. One domain where this extension is particularly relevant is binocular vision. Coordinated activity across both eyes is essential for maintaining single, stable visual perception, supporting everyday activities such as reading and interacting with digital screens. Although normally functioning binocular vision demands little effort operating automatically, impairments in this system can result in significant functional limitations ^40^. Among the core mechanisms of binocular vision is vergence - the inward or outward rotation of both eyes to fixate on objects. Sustained convergence (inward eye rotation) is required for visually near work such as reading.

Despite the central role of convergence in binocular vision and oculomotor control, the network-level mechanisms that generate convergence-related activations remain poorly understood. Several cortical regions have been consistently implicated in the control of convergent eye movements, including the frontal eye fields, supplementary eye fields, parietal eye fields, and primary visual cortex ^41–48^. While these findings underscore the role of localized processing in binocular motor responses and have identified region-specific contributions, it remains unclear whether convergence emerges primarily through localized computations or through interactions across distributed functional networks. If distributed processes do contribute to convergence control, key questions arise: how do known convergence-related regions interact across whole-brain networks, and which specific systems play dominant roles in shaping this functional output? Clarifying this distinction is critical for understanding how large-scale brain systems coordinate precise motor behavior and may offer insight into the neural basis of convergence dysfunctions in both typical visual coordination and its disruption in clinical populations, such as those with convergence insufficiency, concussion-related oculomotor dysfunction, or attention-related disorders.

To address this gap, we tested the hypothesis that convergence-related task activations can be generated via resting-state functional connectivity (FC) using a distributed activity flow mapping framework. Specifically, we hypothesized that functional activations within the regions associated with convergent eye moments are not solely generated through local and regional computations, but also through how these regions are connected to other brain regions through networks, which shape their activations by propagation of activation across large-scale brain-wide networks. Our goal is to investigate how specific brain regions respond to convergent eye movements, resulting from the flow of activations over these regions’ FC through large-scale resting-state networks. This approach builds on prior evidence that intrinsic connectivity patterns can account for localized responses across both perceptual and motor domains.

Given that effective binocular vision requires precise and coordinated input from both eyes, we modeled this process using fMRI data from healthy participants, allowing us to characterize the large-scale network architecture supporting this core oculomotor function. Using resting-state and task-evoked fMRI data from binocularly and neurologically normal (healthy) participants, we investigated a defined set of regions comprising the Cortical Circuitry of Convergence (CCC), encompassing visual and oculomotor areas involved in binocular convergent eye movements. For each region in the CCC, we quantified both empirical task-evoked activation and model-based activation generated by activity flow mapping based on empirical resting-state FC. This framework enabled us to evaluate the extent to which whole-brain, distributed interactions contribute to convergence-specific responses. To further assess the robustness of this network-based model, we leveraged a subset of participants from the same cohort who underwent repeat scanning for resting-state. Building on the assumption that resting-state functional networks remain consistent in healthy individuals, we investigated whether connectivity patterns derived from one session could reliably generate convergence-related task activations in a separate session from the same individuals. This also allowed us to evaluate the reproducibility of activity flow generations and the extent to which intrinsic connectivity patterns can support consistent functional responses.

We propose that functional brain activity in regions associated with convergent eye movements is processed not only through local computations within these areas but also by characteristics of the distributed network-level functional structure of the brain. Specifically, we propose that this includes the FC signatures of these regions. Our results demonstrate that convergence-related functional activations are primarily shaped by resting-state network topology, with early visual and dorsal attention networks contributing most prominently to distributed activations within the CCC. Critically, we observed consistent model accuracy in a subset of participants with repeated resting-state fMRI scans, reinforcing the robustness of this distributed network model. Taken together, our findings suggest that binocular convergent eye movements are governed by the brain’s large-scale functional network architecture, wherein convergence-related instructions propagate through connectivity patterns already present during the resting-state.

## Results

We present our results in five sections. First, we report the overall accuracy of the model used in this study to examine the distributed processes associated with convergent eye movements. Second, we focus on specific brain regions to demonstrate that local responses related to convergent eye movements are reflected in the propagation of functional activations across resting-state FC. Third, we identify which distributed brain networks play a more significant role in generating local activations associated with convergent eye movements. Fourth, we report the full results from the repeated resting-state scan. Finally, we provide results from constrained submodels to determine whether models with limited networks can account for convergence-related activations.

### Distributed network-level processes in shaping local activations

To test our hypothesis regarding the involvement of brain networks in generating the local activations associated with convergent eye movements, we employed approaches to strengthen the inferential power of our generative model. We first began by establishing a strong basis that represents the underlying connectivity structure of the brain to test the mechanisms at the distributed network-level involved in emerging convergent eye movements.

To do this, we used the whole cortex-wide functionally-defined parcellations, known as the Glasser surface-based cortical multimodal parcellation (MMP) atlas ^49^ to construct the functionally connected structure of the brain (Figure 1A). To assign each parcel of the Glasser MMP atlas to a functional network, we further utilized the Cole-Anticevic brain-wide network partition (CAB-NP, Figure 1A). Using a Louvain community detection algorithm on the resting-state fMRI data from the Human Connectome Project, the CAB-NP atlas assigns each region of the Glasser MMP to 12 functional networks ^50^.

**Figure 1.**
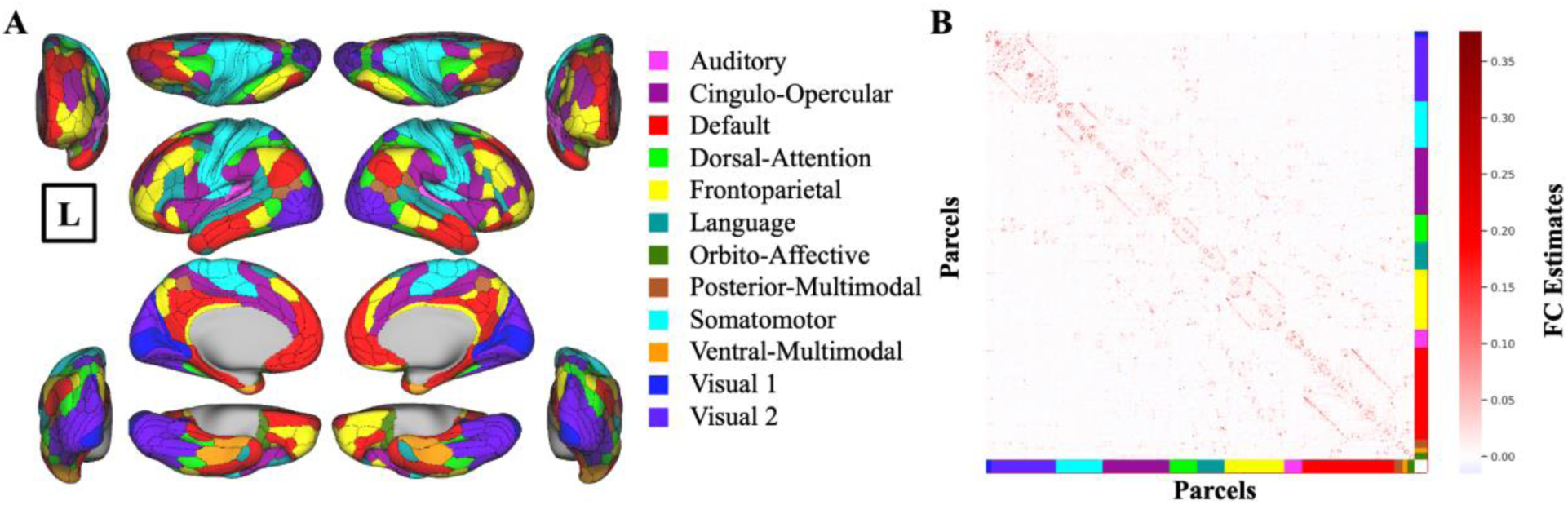
Functional connectivity estimates used for generating activations within the Cortical Circuitry of Convergence. **A:** Glasser MMP atlas illustrated with CAB-NP network assignments. Colors in each parcel represent the network names shown on the right. Brain figures are shown in all planes, Medial-Lateral, Anterior-Posterior, and Dorsal-Ventral. **B:** Group-average FC estimates with regularized partial correlation, first calculated for each participant, and then averaged across all participants. Supplementary Figure S1 shows FC estimates with field-standard Pearson correlation and single-participant FC estimates for a sample participant. Connectome Workbench was used to create all brain figures. “L” sign denotes the left hemisphere.

Second, to avoid the probable inflated results obtained from the field standard Pearson correlation in estimating the number of connected regions ^51,52^, we employed the recently validated and reliable regularized partial correlation method to construct the FC ^51^, which effectively accounts for various widespread confounding factors (Figure 1B).

Third, we used the concept of foundational resting-state fMRI data and used a repeat resting-state fMRI dataset from a subsample of participants for this analysis. Our goal was to re-examine whether the structure of resting-state FC could consistently generate the same local brain activations when the participants were scanned a second time (see below for results, section Repeated Resting-State Scan).

Finally, to ensure that the activity flow-generated activation for each region was not influenced by other functionally related regions associated with convergent eye movements, we excluded all these regions from the source set when computing the activity flow-generated activations (see below for results, section activity propagation).

Before focusing specifically on regions associated with convergent eye movements, we first validated our model using resting-state FC and task-evoked activations from eight task conditions, including the main (convergence-related) and control task conditions. These included two convergence-related tasks (Vergence Motor task (VM) and Vergence Sensory (VS) task), an additional eye movement task unrelated to convergence (Saccadic Motor task), and one motor task of finger tapping. Note that each of the four tasks had two task conditions, resulting in eight task conditions in total (see Methods for more information). To evaluate overall model performance, we compared activity flow-generated and actual brain activation patterns across all conditions and brain parcels (360 cortical parcels from the Glasser MMP atlas). Activity flow-generated activations were computed by weighting the actual activations of source regions by their FC with each target region (see Methods). Model accuracy was quantified as the similarity between activity flow-generated and actual activation patterns, and results were averaged across participants to assess consistency at the group level. This yielded a high overall model accuracy, as indicated by a Pearson’s correlation coefficient (r) of 0.69, an R² of 0.47, and a mean absolute error (MAE) of 0.90. More information on the comparison metrics for model accuracy is provided in the Methods section. Supplementary Figure S2 presents the visualization for the cross-participant comparison of the entire model.

Since our primary focus is the VM task, to assess whether convergence-related activation patterns can be generated across the whole cortex—not just within the brain regions associated with the convergent eye movements—we conducted a benchmark analysis restricted to the VM task. Specifically, we compared activity flow-generated versus actual activation patterns across all cortical parcels during the VM task conditions, across participants. The results indicated substantial correspondence: Pearson’s r = 0.66, R² = 0.44, and MAE = 0.45. This suggests that resting-state FC captures meaningful aspects of task-evoked convergence responses even at the whole-cortex scale. Figure 2 shows the activity flow-generated and actual activation maps for the VM task. Supplementary Figure S3 shows the activity flow-generated and actual activation maps for all task conditions.

**Figure 2.**
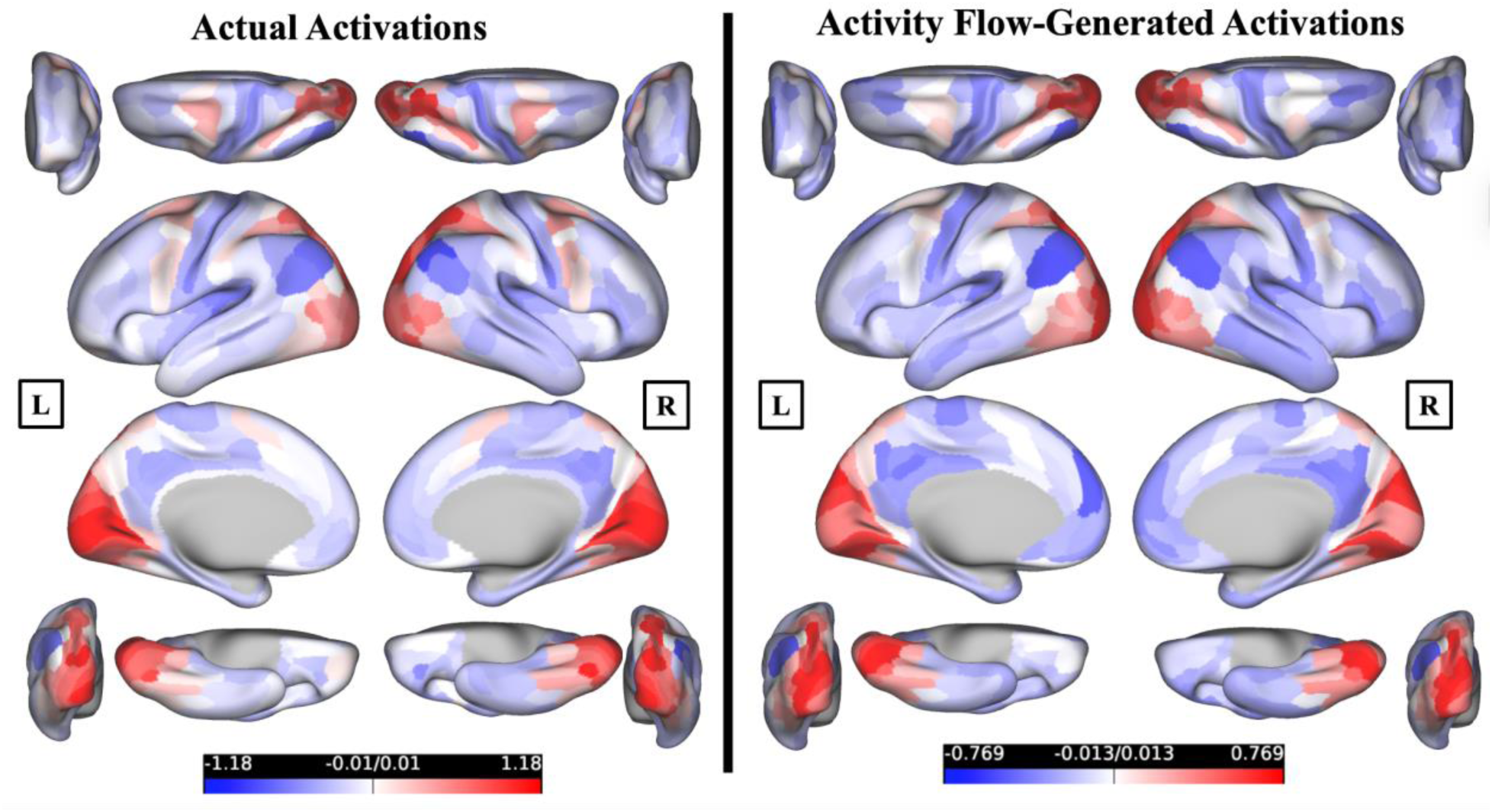
Cross-cortex actual and activity flow-generated activations for the VM task. Actual activations (β values) for the VM task are shown in the left brain plots, averaged across all participants (n = 48). Activity flow-generated activations (β) for the VM task are shown in the right brain plots, averaged across all participants (n = 48). Supplementary figure S3 shows the activity flow-generated and actual activation maps for all task conditions. Connectome Workbench was used to create all brain figures. “L” and “R” signs denote left and right hemispheres, respectively.

### Activity propagation over resting-state functional connectivity generates convergence-evoked activations in oculomotor regions

By utilizing the hypothesis of distributed processing in the convergence domain, we evaluated whether convergence-specific responses within the oculomotor regions are primarily shaped by activity flowing through their intrinsic resting-state FC with other brain regions. This could show that these regions’ unique connection to all other regions may shape their task-evoked responses during a convergence task. We refined the analysis to focus specifically on these regions to extend the whole-cortex evaluation and test our model more directly on the brain regions involved in convergent eye movements. We selected brain regions responsible for convergent eye movements in both hemispheres, referring to this functional circuitry as the “Cortical Circuitry of Convergence (CCC)” in both hemispheres (see Methods, Figure 3A), and computed their unique FC (see Methods, Figure 3B). This refined analysis focused on the CCC across all eight task conditions—collectively referred to as the response profile or population receptive field of these regions—in both hemispheres. This targeted evaluation allowed us to confirm the responsiveness of the CCC to convergence-specific eye movements, while also assessing the model’s generative validity across a broader set of other eye movement and motor learning tasks. This step served as an additional confirmation of the model’s generalizability beyond convergence-specific activation alone. Regions of interest within the CCC were defined a priori based on previous neuroimaging literature ^41–47,53–56^ as well as repeatability analysis of convergent eye movements ^48^. These selected brain regions span beyond the visual cortices and include areas across the cortex. Specifically, four regions defined in each hemisphere were: the primary visual cortex, the frontal eye fields, the supplementary eye fields, and the parietal eye fields. These regions map onto the Glasser MMP atlas within three major resting-state networks as defined by the CAB-NP: Visual 1 (primary visual cortex), Cingulo-Opercular (the frontal eye fields and the supplementary eye fields), and Dorsal Attention (parietal eye fields). Although the anatomical distribution of these regions of interest (ROIs) from early visual cortex to frontoparietal control systems suggests a role for distributed processing, our analysis formally tested this hypothesis. Note that when computing the activity flow-generated activations in each of the eight regions within the CCC in either hemisphere, we excluded all other CCC regions from the source set.

**Figure 3.**
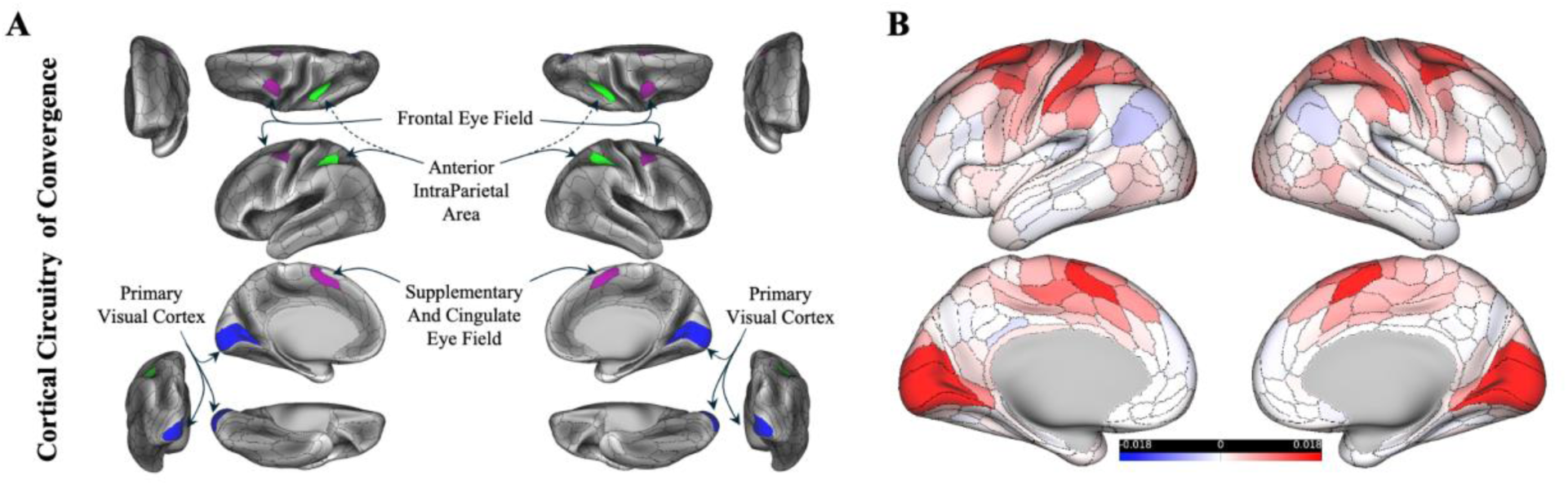
Cortical Circuitry of Convergence. **A:** Parcels associated with the convergent eye movements in both hemispheres. Colors in each parcel match those shown in Figure 1A, specifically with CAB-NP. **B:** Resting-state FC estimates using graphic lasso seeded to the Cortical Circuitry of Convergence. Connectome Workbench was used to create all brain figures.

Both hemispheres demonstrated high mapping accuracy in the response profile. In the left CCC, the activity flow-generated versus actual activation profile yielded a Pearson correlation coefficient of *r* = 0.86, *R²* = 0.52, MAE = 0.6. The results of the right CCC were similarly strong (*r* = 0.77, *R²* = 0.37, and MAE = 0.67). Building on prior findings across various cognitive tasks ^35^, our results confirm that the model accurately generates activation within the CCC across a range of visual and non-visual conditions. By leveraging the resting-state FC of these regions with the rest of the brain, the findings underscore the role of distributed network-level processes in generating localized activations within the oculomotor convergence system.

Additionally, a critical question is whether the regions within the CCC are sensitive to the motor response part of the convergent eye movement (efference pathway), rather than simply observing the task stimulus or the sensory portion of the circuit (afferent pathway). For this reason, we refined the previous benchmarking analysis for the VM task by focusing on the CCC, in both actual and activity flow-generated activations. If these regions are specifically responsible and associated with the active convergent eye movements, this could be seen in both actual and activity flow-generated activations. We tested this question in two ways. First, we tested whether these regions are significantly responsive to active convergence step demands, rather than general visual input. While the previous benchmarking established that distributed resting-state FC can approximate whole-cortex activation during convergence motor execution (VM), this analysis aimed to determine whether the activity flow-generated activation patterns within the CCC were actively engaged in the motor response of the convergence demand. We contrasted activation levels in the VM task with those from a sensory control task (VS), in which participants passively viewed the same stimuli without performing convergence step responses. Significant differences emerged in both actual and activity flow-generated activations, with greater activation in VM relative to VS in both hemispheres (all p < 0.00001, corrected for multiple comparisons using non-parametric permutation testing with the max-T method; see Methods). In the actual data, t(47) = 4.85 for the left hemisphere and t(47) = 6.98 for the right hemisphere (Figure 4A and 4E). In the activity flow-generated data, t(47) = 3.7 for the left hemisphere and t(47) = 3.37 for the right hemisphere (Figure 4B and 4F). These results confirm that the CCC exhibits selective engagement during motor execution of convergence, and that this specificity is generated from the flow of activations through distributed resting-state FC patterns.

**Figure 4.**
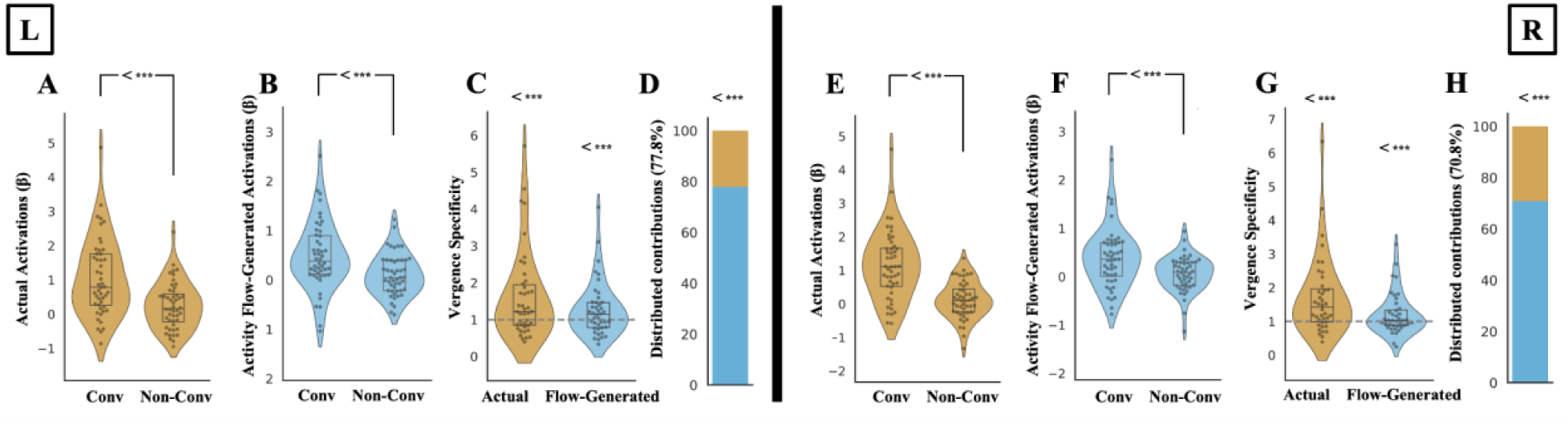
Distributed processes in the cortical circuitry of convergence are sensitive to the motor response of the convergent eye movement. **A&E.** Actual activations in the VM and VS tasks in the left and right CCC, respectively. **B&F.** Activity flow-generated activations in the VM and VS tasks in the left and right CCC, respectively. “Conv” denotes convergence-related oculomotor task conditions (VM task) that respond to the motor demand of the task execution, and “Non-Conv” denotes task conditions related to the sensory demand of the task execution (VS task). **C&G.** Vergence specificity calculated for actual and activity flow-generated activations in the left and right CCC, respectively. The dashed grey line indicates vergence specificity null hypotheses of 1, as VM activations are similar to the VS activations. **D&H.** Distributed contribution percentages in the left and right CCC, respectively. *** Denote significant results from the upper-tailed paired or one-sample t-test using nonparametric permutation testing with 100000 permutations, corrected for multiple comparisons with Max-T.

Although these results indicate that the CCC is significantly more responsive to active motor execution compared to passive visual stimulation in both hemispheres, the responses observed during the sensory stimulation only (VS task) could still be relatively high, even though they did not significantly reach the levels seen in the voluntary motor task (VM task). To further investigate this issue, we calculated a “vergence specificity index” in the next step of our analysis pipeline. After normalizing activation values using a standard min-max normalization (see Methods) to ensure interpretability and comparability across participants by eliminating negative values and rescaling all activation values to a common [0, 1] range, we calculated the vergence specificity index as the ratio of activations during the VM task to activations during the VS task, separately for actual and activity flow-generated responses within the CCC. This index demonstrated the specificity of the regions within the CCC in terms of actually performing convergent eye movements compared to passively observing the task stimuli. The ratio of 1 in this index shows similar responses in VM and VS tasks in both actual and activity flow-generated activations. Nonparametric permutation testing with 100,000 permutations for both actual and activity flow-generated activations revealed statistically significant vergence specificity in the left and right CCC, as compared to the null hypothesis of VM activations being similar to VS activations (null hypothesis of ratio as 1, p < 0.00001, corrected for multiple comparisons with max-T). Significant t-values for the left hemisphere were t(41) = 3.38 for the actual and t(41) = 2.36 for the activity flow-generated activations (Figure 4C). Right hemisphere values were t(39) = 3.86 for the actual and t(39) = 2.11 for the activity flow-generated activations (Figure 4G). The degrees of freedom in this context are determined by the sample size, with the exclusion of any outlier participants. Outlier removal (see Methods) in the vergence specificity index excluded a small number of participants in both hemispheres whose vergence specificity scores were not representative of the group average. Running all computations without removing outliers showed no change in the statistical significance level of the results.

Furthermore, the ratio of activity flow-generated over actual vergence specificity index could be a measure of how the distributed network mechanism manages activations in the CCC. This is because the difference between actual and activity flow-generated activations lies in their underlying mechanisms. Actual activations can result from both local and distributed processes, as well as unpredictable noise that influences convergence step responses. In contrast, activity flow-generated activations are solely based on a model that considers the interactions between FC and brain-wide activations. Therefore, this ratio reflects the actual activations that can be explained by the distributed network mechanisms leading to activity flow-generated activations. Both hemispheres showed statistically significantly higher ratios than 50% (p < 0.00001, corrected for multiple comparisons with max-T). Distributed network-level contributions to the vergence specificity in the CCC were 77.85%, t(41) = 7.1, in the left hemisphere (Figure 4D) and 70.84%, t(39) = 5.82, in the right hemisphere (Figure 4H). Similarly, repeating computations without removing outliers showed no change in the statistical significance of the results.

### Quantifying the contributions of large-scale resting-state networks to local convergence-evoked activations

Our hypothesis regarding the network-level distributed process was based on two key observations. First, the regions associated with convergence generation, as reported in earlier studies, are actually dispersed across various networks throughout the cortex (see Results above and Methods). Second, since these regions are not adjacent to one another, it primarily suggests that certain networks may play a more significant role in the emergence of the convergence-related response within the CCC. Therefore, we aimed to find which of the 12 cortical networks (as defined by CAB-NP) contributes most significantly to this process (see Methods). Based on their CAB-NP network assignment, an average of the “flows” (propagations) directed toward the CCC could provide a network-level mapping of flows to the CCC. Further comparisons could reveal which networks have a greater impact on the activations of the CCC. Comparisons were evaluated using a max-T threshold of t = 3.45 (p < 0.00001) for the left hemisphere and t = 3.43 (p < 0.00001) for the right hemisphere, based on 100,000 permutations. Significance was determined by whether the observed t-statistic exceeds the corresponding max-T threshold. The results indicated that VIS2 and DAN significantly contribute the most to the activations in the CCC, compared to other networks. Specifically, the contributions rank as follows: VIS2 > DAN > all other networks, based on the number of significant comparisons out of 11 comparisons to all other networks. We observed this pattern in both hemispheres, with statistical significance noted (p < 0.00001, corrected for multiple comparisons using the maximum T method). Figure 5A and 5B show the ranked bar plots representing the total contribution strength of each network (i.e., average t-values). Supplementary Figure 4 shows all network-to-network comparisons for both hemispheres along with their observed t-statistics.

**Figure 5.**
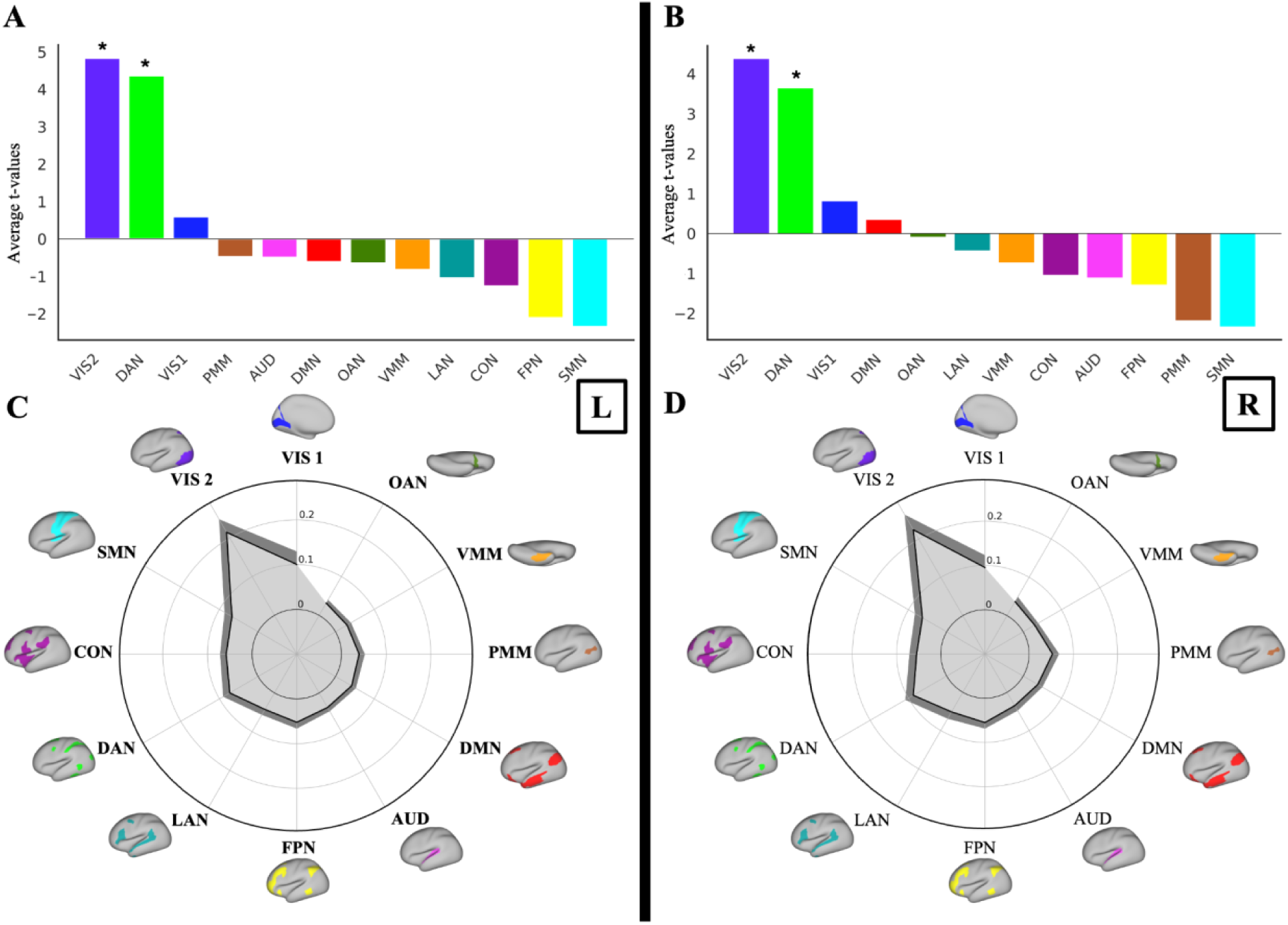
Contributions of large-scale resting-state networks to local convergence-evoked activations. **A&B**: Ranked bar plots showing the total contribution strength of each network (i.e., average t-values) to activity flow generation. VIS2, and DAN followed by VIS1, consistently ranked highest in left and right CCC in the VM task. Asterisks show two significant networks. **C&D**. The variance explained by each network-restricted activity flow model (unmixed partial R² shown in gray color) of the left and right CCC response profile. Dark grey shows 95% confidence interval across participants. Similarly, VIS2, VIS1, and the DAN were ranked as the first three networks in the dominance analysis in both hemispheres.

We then calculated the contributions of each network at the response profile level using dominance analysis. We assessed the unique contributions (partial R²) of each of the 12 networkers to the activity flow-generated response profile of the CCC by recomputing the activity flow-generated activations across all task conditions while restricting source regions to their CAB-NP network assignments (see Methods). This approach effectively parsed the entire model’s generated activations into network components, ensuring that the sum of all these network-level values equaled the overall model value (see Methods). Dominance analysis further identified VIS2, VIS1, and the DAN as the most influential networks in accounting for variance within the CCC in both hemispheres (Figures 5C and D), highlighting the contribution of low-level visual and attentional networks in driving the convergence-related activation profile. We report values for all networks in Supplementary Table 1. Overall, across both the conditions explicitly associated with the convergence motor task and the full set of task conditions, including both convergence-related and non-convergence-related tasks, visual and dorsal attention networks consistently emerged as the most influential contributors to activations within the CCC.

We also probed and visualized these activity “flows” into the left and right CCC under VM (oculomotor vergence) and VS (sensory only) task conditions (Figure 6). Note that each activation generated by activity flow mapping is computed as the sum of individual flow terms, defined as the product of a source region’s activation and its resting-state FC estimate with the target region. These flow terms represent the distributed contributions across the brain that result in the generated activation in a region. To better understand the specificity of convergence-related responses, we examined how these individual flow terms differed between the convergence motor task (VM) and the sensory task condition (VS). This allowed us to identify the distributed sources that most strongly drive task-related activation in the CCC and to assess whether distinct patterns of distributed network interactions support motor-related demands. To statistically assess differences, we conducted a paired, upper-tailed max-T permutation test (100,000 permutations) across subjects for each network. The results revealed significantly greater flow terms within the VIS2 network during VM compared to VS. To examine the spatial resolution of these effects, we extended the analysis to the parcel level. For each of the 360 Glasser parcels, we computed subject-wise differences in flows between VM and VS, then applied a max-T permutation test to identify parcels with significantly stronger flow during VM. Although the group-average maps showed prominent visual differences in multiple parcels, only a limited number of parcels survived correction for multiple comparisons. A subset of parcels exhibited significantly greater flow during VM, predominantly located within the VIS2.

**Figure 6.**
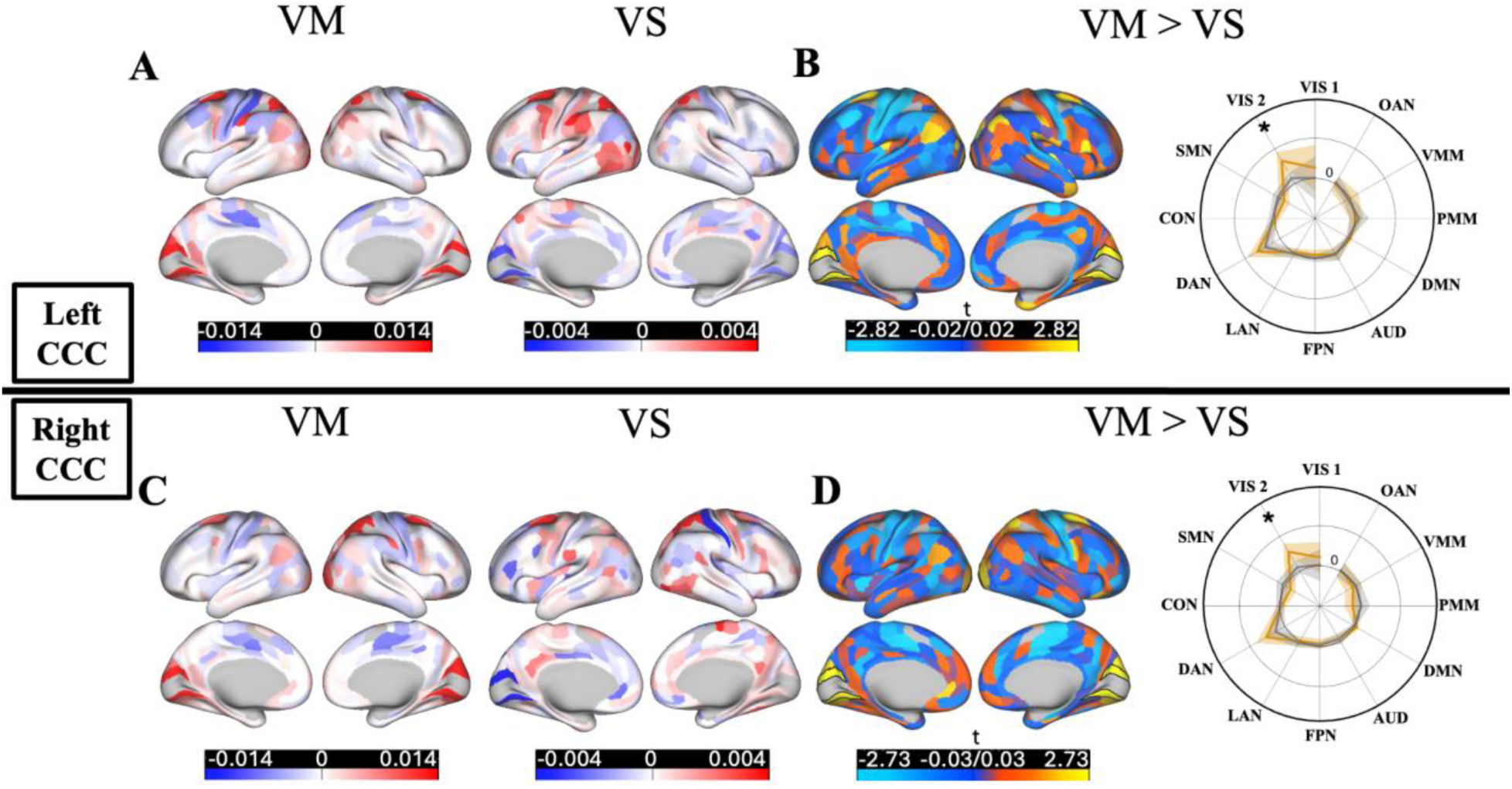
Activity flows in VM and VS task conditions. **A&C.** Brain-wide activity flow terms into the left (top row) and right (bottom row) CCC during VM (first column) and VS (second column) task conditions. **B&D.** VM>VS contrast in the flow terms, contrasted in the parcel level, as well as the average-network level into the left CCC (top row) and right CCC (bottom row). Parcels with significantly greater VM > VS flows are outlined in black borders. Polar plots show the mean activity flow from each CAB-NP cortical network into the CCC, with shaded regions indicating 95% confidence intervals across subjects. Network-level activity flows were significantly greater during VM than VS within the VIS2 network (p < 0.00001, Max-T corrected across 12 networks). Note that eight regions within the left and right CCC did not contribute to generating the activations, hence shown in grey color in all brain plots.

### Repeated Resting-State Scan

Of the 48 participants included in the primary analysis, 27 underwent a repeat session using identical protocols within 1–31 days (mean 7.67 ± 7.72 days) after the initial scan, based on the availability of the participant and the imaging center. This provided a unique opportunity to evaluate whether resting-state FC serves as a reliable and individualized signature that can generate task-evoked activations. For each of these 27 participants, we used resting-state data from the second scan to model task-evoked activation patterns observed in the initial session. This approach directly tested our hypothesis that activation within the CCC can be generated through a distributed activity in other brain regions, weighted by their unique resting-state FC to these regions. Given the known stability of resting-state FC across sessions in healthy individuals ^57^, this analysis further validated the predictive utility of distributed network interactions. In addition to overall model accuracy, we extended this analysis to examine the whole cortex VM task, response profile, vergence specificity, and network contribution.

Our results showed a high overall model accuracy (Pearson’s correlation coefficient r of 0.70, an R² of 0.48, and MAE of 0.92) and a similarly substantial correspondence in the whole cortex mapping accuracy in the VM task (Pearson’s r of 0.70, an R² of 0.48, and MAE of 0.92). Both CCCs showed high accuracy in the response profile in the left hemisphere (Pearson’s r of 0.86, an R² of 0.45, and MAE of 0.66) and the right hemisphere (Pearson’s r of 0.81, an R² of 0.34, and MAE of 0.67). Vergence specificity results in the left CCC were t(23) = 2.72 for the actual and t(23) = 1.34 for the activity flow-generated activations (Figure 7); and in the right CCC were t(20) = 3.11 for the actual and t(20) = 1.36 for the activity flow-generated activations (Figure 7). Distributed network-level contributions to the vergence specificity in the left CCC were 74.86%, t(23) = 4.93, (Figure 4D) and 75.97%, t(20) = 5.23, in the right CCC (Figure 7). Similarly, repeating computations without removing outliers showed no change in the statistical significance of the results.

**Figure 7.**
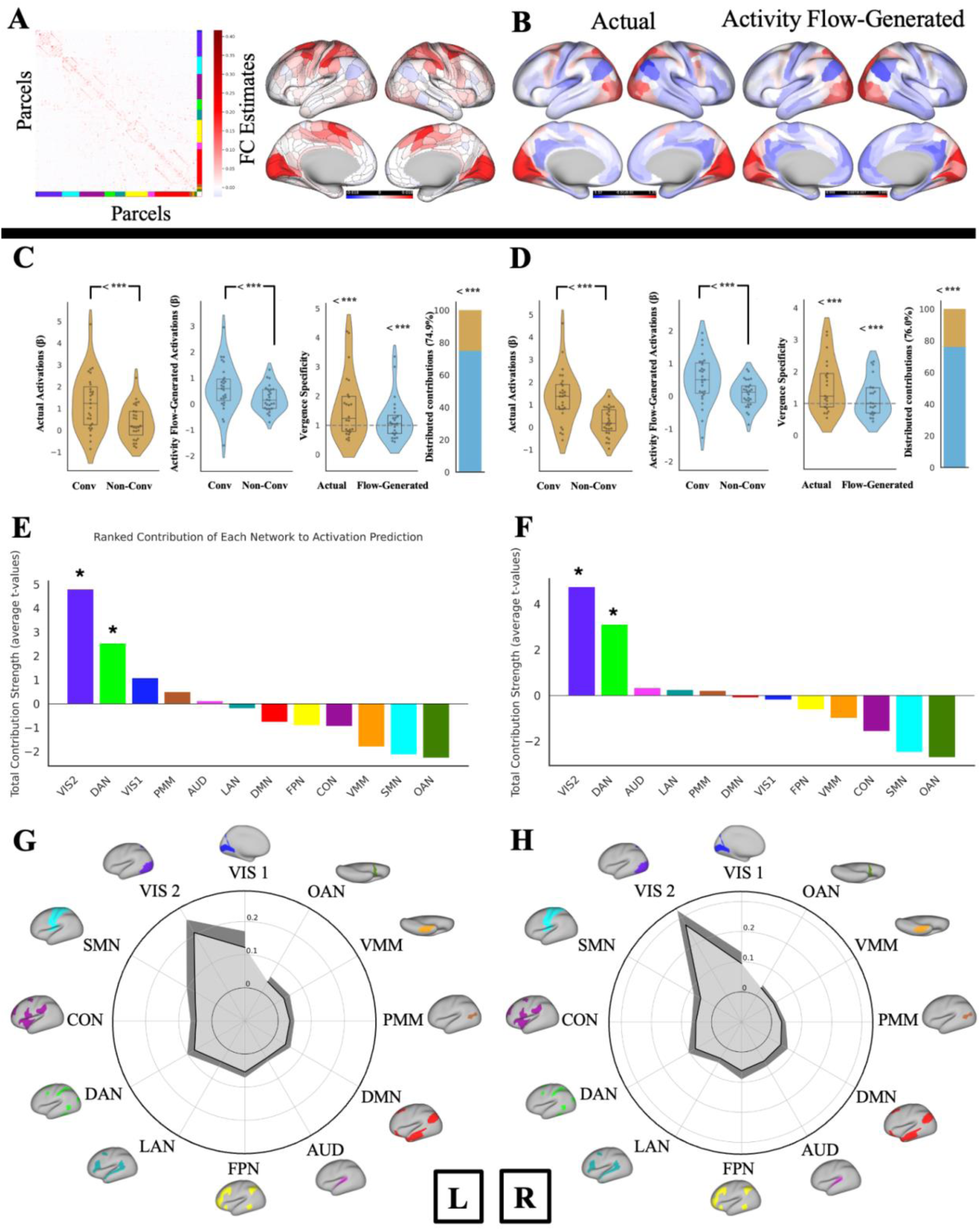
Overall results with Repeated Resting-State Scan. **A.** Functional connectivity estimates from the repeated resting-state scan used for generating activations within the Cortical Circuitry of Convergence (CCC). Group-average FC estimates with regularized partial correlation, first calculated for each participant, and then averaged across all participants. The brain plot shows the resting-state FC estimates from the repeat scan using the graphic lasso seeded to the Cortical Circuitry of Convergence. **B.** Cross-cortex actual and activity flow-generated activations (β) for the VM task conditions (similar to the primary analysis). **C-D.** Actual and Activity flow-generated activations in the VM and VS tasks. Vergence specificity was calculated for actual and activity flow-generated activations, and distributed contribution percentages in the left and right CCC, respectively. “Conv” denotes convergence-related oculomotor task conditions (VM task) that respond to the motor demand of the task execution, and “Non-Conv” denotes task conditions related to the sensory demand of the task execution (VS task). The dashed grey line in the vergence specificity graphs indicates the vergence specificity null hypothesis of 1, as VM activations are similar to the VS activations. *** Denote significant results from the upper-tailed paired or one-sample t-test using nonparametric permutation testing with 100000 permutations, corrected for multiple comparisons with Max-T. **E.** Ranked bar plots showing the total contribution strength of each network (i.e., average t-values) in the left hemisphere. **F.** Ranked bar plots showing the total contribution strength of each network (i.e., average t-values) in the right hemisphere. **G.** The variance explained by each network-restricted activity flow model (unmixed partial R² shown in gray color) of the left CCC response profile. **H.** The variance explained by each network restricted activity flow model (unmixed partial R² shown in gray color) of the right CCC response profile. The dark grey represents a 95% confidence interval across participants.

The results of the network contributions were also consistent with the primary analysis. The results in both hemispheres indicated that the VIS2, and the DAN contribute the most to the activations in the CCC, compared to other networks, as VIS2 > DAN > all other networks, based on the number of significant comparisons out of 11 comparisons to all other networks (p < 0.00001, corrected for multiple comparisons using the maximum T method). Supplementary Figure 5 presents all network-to-network comparisons for both hemispheres, along with their corresponding observed t-statistics. Dominance analysis further identified VIS2, VIS1, and the DAN as the most influential networks in accounting for variance within the CCC in both hemispheres (Figure 7), highlighting the contribution of low-level visual and attentional networks in driving the convergence-related activation profile. We report values for all networks in Supplementary Table 1. Overall, across both the conditions explicitly associated with the convergence motor task and the complete set of task conditions (including both convergence-related and non-convergence-related tasks), visual and dorsal attention networks consistently emerged as the most influential contributors to activation within the CCC.

### Control Analyses for Circularity Concerns and Potential Inferential Limitations

To address potential concerns about circularity in our analysis, we implemented several methodological controls to ensure that activity flow estimates were not biased by information leakage from functionally related regions. The most critical of these controls involved how the regions within CCC were handled during activity flow mapping. Fundamentally, the activity flow mapping framework estimates the activation of each region by holding it out and computing the weighted flow of activations from all other regions in the brain. To more rigorously prevent circularity in generating convergence-related activation, all our analyses included a conservative and rigorous step. When computing the activity flow-generated activations in each of the eight regions within the CCC in either hemisphere, we excluded all other CCC regions from the source set. This ensured that the activity flow-generated activation for each region was not influenced by other functionally related regions within the right and left CCC; regions that play roles in convergent eye movements. Notably, a version of the model that retained the rest of the CCC (excluding only the held-out region) showed overall similar model accuracy (Pearson’s r of 0.70, an R² of 0.48, and MAE of 0.89). However, to maintain strict control for regional interdependence, all results reported throughout the manuscript regarding the whole cortex distributed mechanisms are based on the model in which all eight CCC regions were excluded from activity flow mapping.

To test whether spatial smoothness-induced circularity was driving our activity flow results, in a control analysis we excluded all parcels within 10 mm of each target region, which was additional to holding out the eight predefined convergence-related ROIs from contributing to each other’s mappings. Activity flow-generated results remained robust after these exclusions, with a mean correlation between activity flow-generated and actual task activations of Pearson’s r = 0.54, (one-sample t test versus zero, t(47) = 33.64, p < 0.00001), confirming that the observed activity flow effects were not merely a byproduct of spatial smoothness or ROIs interdependence, but instead reflect meaningful functional interactions captured by resting-state FC.

#### Condensed Cortical Models

##### No additional benefits of circulating the signal with the CCC to improve the generated activations

To explore whether reciprocal or recurrent processes (as proposed by ^58^) within the convergence system contribute to activation dynamics, we extended our original activity flow model to include potential bidirectional signal flow within the CCC. While our primary analysis excluded all CCC regions when mapping activation within them, this follow-up analysis aimed to test whether internal circulation among CCC regions could further improve activity flow-generated activations and better capture the representational dynamics of convergence-related activity.

Specifically, we hypothesized that once distributed activity (flows) reaches to the CCC, a subsequent step of intra-circuitry flow (representing potential reciprocal interactions) might refine the activity flow-generated activation patterns. To test this, we used the initial activity flow-generated activations (from the primary model) as inputs and computed a secondary round of activity flow within the CCC. As before, each region was held out in turn, and its activation was generated using the remaining seven regions and their resting-state functional connectivity. Note that holding out all the CCC regions from the source regions was theoretically not possible here because the signal was allowed to circulate only within these eight regions. Therefore, only the target region (to be computed) was held out each time. This approach generated a new set of activity flow-generated activations that reflected one step of potential internal circulation within the right and left CCC. Whether additional rounds of within-CCC propagation were necessary or not depended on whether this secondary activity flow-generated activations improved model performance. Results showed that while the secondary generated activations preserved significant vergence specificity in the generated activations in both hemispheres (t(39) = 1.94 for the left hemisphere and t(41) = 2.54 for the right hemisphere, p < 0.00001, corrected for multiple comparisons with max-T), they did not improve upon the primary activity flow-generated activations. Notably, *R²* values dropped substantially in the secondary circulation from 0.52 to - 0.26 in the left and 0.37 to −0.25 in the right hemisphere, indicating that the secondary activity flow-generated activations could not validly explain variance in the actual activation patterns (Figure 8). These findings suggest that once activities from distributed cortical networks reach the CCC, these flows are sufficient in generating activations within the CCC, and further internal propagation among its regions (internal dynamics) does not add unique explanatory power concerning task-evoked responses.

**Figure 8.**
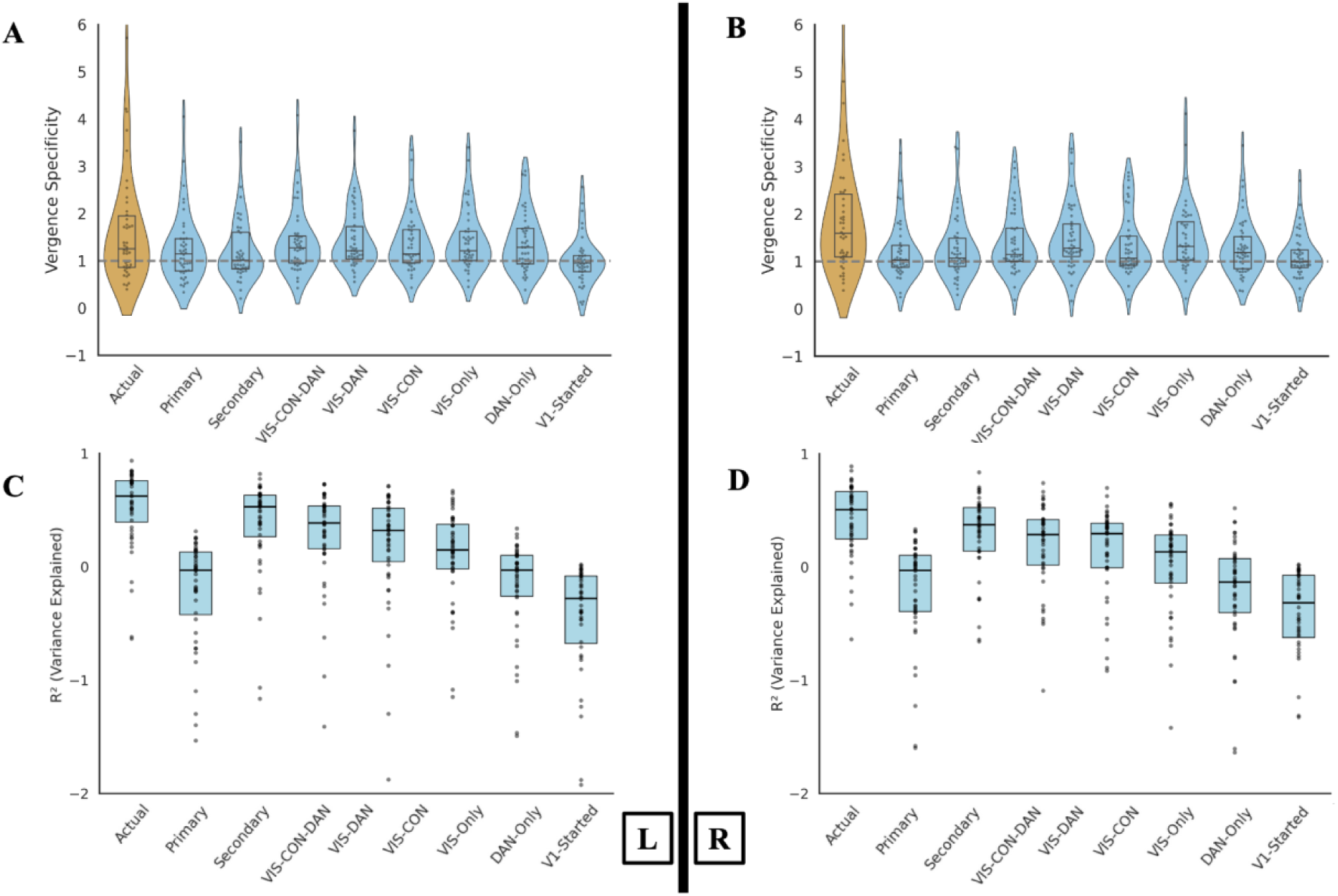
Results of Submodel. **A–B.** Vergence specificity of actual and activity flow-generated activations in the left (A) and right (B) CCC under various activity flow constraints. **C–D.** Activity flow mapping accuracy (*R²*, variance explained) in the left (C) and right (D) CCC across the same constraint conditions.

##### Better results of the whole-cortex distributed processes compared to condensed cortical submodels

We next refined our analysis by condensing the networks contributing to the distributed activity flow mapping. This hypothesis was tested to ensure the robustness of the whole cortex distributed process in generating activations within the CCC. Additionally, this could answer the question of whether each of these models (with limiter networks) is sufficient to generate vergence specificity and response profiles in the CCC. If so, then the inference about the role of the whole cortex distributed process in generating vergence specificity and activations with CCC could be discussed in relation to these networks only. For example, since one of the regions of the CCC is within the DAN, the combination of the Visual and DAN networks could be sufficient for generating activations within the CCC. Therefore, a combination of these hypotheses was available to test. To do this, we implemented several submodels, each consisting of only one or a limited number of networks. Since the regions with the CCC are already part of the Visual, Dorsal Attention, and Cingulo-Opercular networks, the first model included brain parcels within the visual networks (VIS-1 and VIS-2), CON, and DAN (VIS-CON-DAN). Other models included Visual networks and CON (VIS-CON), Visual networks and DAN (VIS-DAN), and an individual network as Visual networks only (VIS-Only). Since DAN significantly and consistently interacted with the network contribution in the primary analysis, this particular network was also tested in a separate model (DAN-Only) as an exploratory submodel. However, all these models were control models meant to refine inferences relative to the primary whole-cortex distributed model, including all 12 networks. Collectively, all these models and possibilities could confirm whether a refined model is sufficient to generate activations within the CCC, or whether the whole-cortex distributed mode generates activations most accurately, although with potential inferential limitations. We further focused on a model in which signals could only be initiated from V1 and propagate (through their resting-state FC) towards the other regions within CCC (the V1-Initiated model). This model could help rule out the hypothesis that V1-initiated activations are sufficient in generating the vergence-related activations within the CCC.

For each model, we extracted the FC estimates using only the parcels related to the network(s) within the model, along with the CCC regions (if they were outside of these networks). For example, in a model focused on the Visual and DAN networks, we included all regions within the VIS-1, VIS-2, and DAN, as well as four CCC regions (2 in each hemisphere) in the CON. For all models except the V1-Initiated model, all CCC regions were held out when generating the activations in each CCC region. For the V1-initiated model, we used mean-centered activations in the two V1s as source activations to generate the activations of the other six CCC regions. Mean-centered activations in each V1 region were used to compute activations in the other hemisphere’s V1. In all models, the activations in the included source regions, weighted by their FC estimate, shaped the activations in the target regions.

The significant results for the vergence specificity in these submodels for the left hemisphere were VIS-CON-DAN: t(41)=3.72, VIS-CON: t(40)= 3.78, VIS-DAN: t(41)= 4.29, VIS-Only: t(41)= 4.25, and DAN-Only: t(42)= 3.96 (all p < 0.00001, corrected for multiple comparisons using non-parametric permutation testing with the max-T method; see Methods). The significant vergence specificity results of the submodels for the right hemisphere were VIS-CON-DAN: t(43)=3.99, VIS-CON: t(43)= 3.36, VIS-DAN: t(42)= 4.64, VIS-Only: t(43)= 4.38, and DAN-Only: t(44)= 3.19 (all p < 0.00001, corrected for multiple comparisons using non-parametric permutation testing with the max-T method; see Methods). Results of the V1-Initiated mode for the left hemisphere were t(41)= −0.12 (p =1), and for the right hemisphere were t(41)= 1.3 (p < 0.00001, corrected for multiple comparisons using non-parametric permutation testing with the max-T method; see Methods, Figure 8). None of these vergence specificity results were superior or improved compared to the preliminary results from fully distributed whole-cortex processes. Note that in a one-tailed max-T permutation test, significance is determined by the empirical permutation threshold; therefore, t=1.3 was significant in this case.

In addition to the vergence specificity in each submodel, we compared the activity flow-generated and actual activations at the response profile level in each of the left and right CCCs, thereby calculating R² values, iterated over held-out regions, task conditions, and participants. In the left hemisphere, *R²* values dropped from 0.52 in the primarily generated activations to 0.37 in the VIS-CON-DAN submodel, 0.28 in the VIS-DAN submodel, 0.18 in the VIS-CON submodel, 0.06 in the VIS-Only submodel, −0.22 in the DAN-Only submodel, and −0.52 in the V1-Initiated submodel. In the right hemisphere, *R²* values dropped from 0.37 in the primarily generated activations to 0.24 in the VIS-CON-DAN submodel, 0.13 in the VIS-DAN submodel, 0.08 in the VIS-CON submodel, −0.05 in the VIS-Only submodel, −0.26 in the DAN-Only submodel, and −0.51 in the V1-Initiated submodel (Figure 8).

These results demonstrate that a submodel with one or a few networks, or even a model initiated from V1 only, cannot explain the variance in actual activations well and is therefore insufficient to describe the brain mechanisms underlying convergent eye movements. Regarding the positive R² values, the results show that only a combination of two or three networks can generate activations with positive R² values, implying a limited explanation of the variance in the actual activations. However, none of these R² values reach the level of the whole-cortex distributed processes. No individual network or a V1-initiated process was able to explain the variance in the actual activations of the CCC. It is worth noting that models with more source regions can appear more accurate because larger predictor sets capture more variance, and more accurate predictions are expected. Therefore, to compare model performance, an option could be to use information criteria, the Akaike Information Criterion (AIC) and the Bayesian Information Criterion (BIC), to penalize model size. However, given the limited number of task conditions per response profile and models ranging from a few to hundreds of source regions, the complexity penalty would dominate the fit term and mechanically favor the smallest models even when their predictions are poor (for example, the V1-started model showed negative R²). Therefore, we did not use AIC or BIC for inference and instead report R² on the response profiles as the accuracy measure.

Collectively, these findings showed that although some of these submodels generated significant vergence specificity, this was not at the level of the vergence specificity generated through a whole-cortex distributed process, nor was it substantially better than that. This implies that vergence specificity is better predicted when the whole cortex is involved in generating the CCC activations. Additionally, and importantly, comparisons of the generated activations with actual activations revealed poor R² values in these submodels, indicating worse activity flow-generated activations when condensed into a subset of one or a few networks. Interestingly, the V1-initiated models generated the worst activations, exhibiting both insignificant vergence specificity and greater negative R² values compared to the actual activations. This suggests a poor fit in these models. Figure 8 illustrates the vergence specificity, along with the related t-value and R² in each model.

## Discussion

Our findings show that binocular convergent eye movement control is primarily governed by distributed, network-level propagation of functional activations over intrinsic resting-state functional connectivity (FC). Regions within the bilateral *cortical circuitry of convergence (CCC)* showed strong specificity for voluntary convergent eye movements, both in actual and activity flow-generated responses. Furthermore, distributed network-level processes significantly accounted for more than half of the variance in vergence specificity observed in the actual activations, suggesting that these responses emerge from distributed processes rather than isolated local activations. Visual networks (VIS1 and VIS2), along with the dorsal attention network (DAN), played a dominant role in initiating convergence-related activation within the CCC. This network-based contribution was consistent not only for convergence-specific tasks but also across a range of control conditions involving visual and motor learning demands, suggesting that the convergence system is integrated within a broader functional network structure. These results suggest that voluntary convergent eye movements are supported by large-scale interactions between early visual and attention networks, which together propagate functional activations toward convergence-related regions.

Additionally, these effects were confirmed in a repeat test of a subgroup of participants from the initial analysis, further demonstrating the reliability of our findings. In this follow-up analysis, we observed that resting-state FC patterns within the CCC, specifically how these regions connect and interact with other brain areas while not engaged in a convergence task, served as a reproducible signature. This signature not only generated task-evoked responses during convergence-specific tasks but also across a wider range of task conditions and even in a smaller sample. This suggests that one acquisition of resting-state FC can serve as a functional signature for brain regions involved in convergent eye movements, leading to consistent and reproducible convergence-related activations across various tasks. This finding aligns with previous literature, indicating that functional brain networks are not solely influenced by daily variations ^57^. Here, we use the term “functional signature” to describe how specific the FC of these regions is to themselves. This distinction is crucial to avoid confusion between the terms “connectome” and “fingerprint,” as defined by Finn et al. ^59^. Based on this, indeed, two key concepts here reinforce one another in supporting the conclusion that regional activations are shaped by distributed, network-level processes. First, resting-state FC provides a stable and consistent measure of the brain’s intrinsic functional architecture. Second, the activity flow mapping framework used here leverages this architecture by modeling how regions within the CCCs are functionally connected to the rest of the brain. These distributed connections determine the flow of activations that shape task-evoked responses. Together, these findings suggest that just as the global structure o resting-state FC remains consistent across individuals, the unique connectivity patterns/signature—or functional circuitry—of specific regions also remain stable. This consistent connectivity profile likely reflects both functional and underlying structural organization and plays a critical role in shaping how these regions respond during convergent eye movement tasks.

Our results build upon prior work demonstrating that localized brain activations can emerge from distributed activity flow processes shaped by intrinsic connectivity patterns ^35,37–39^. While earlier studies primarily investigated these mechanisms in the context of visual stimulus differentiation, we extended this framework to the oculomotor domain, specifically during binocular convergent eye movements. By focusing on regions within the CCC, we tested whether task-evoked activations related to eye movement control also emerge from distributed network interactions. Consistent with prior findings in the visual system, we observed that network-level processes shaped by resting-state FC largely drove convergence-specific activations. This was confirmed by the strong correspondence between actual activations and those generated via activity flow models based on distributed FC. These results align with theoretical accounts suggesting that such models are particularly effective when global coupling dominates over local processing ^35^. Importantly, distributed processes substantially explained how specifically the CCCs responded to the convergent eye movement demands, explaining more than 50% of the variance in actual activations compared to local-only processes. This finding indicates that global integration via intrinsic functional networks contributes meaningfully to regional-level responses, even in a domain involving oculomotor output. Moreover, the generative strength of distributed models generalized across a variety of task conditions, including those unrelated to convergence, underscoring the robustness of resting-state FC as a platform for diverse functional activations. Across both convergence-related and unrelated tasks, the flow-based models consistently identified two visual networks (VIS1 and VIS2) and the dorsal attention network (DAN) as the primary contributors to emergent activation within the CCC.

Additionally, our findings revealed consistently distributed processing mechanisms across both hemispheres, suggesting a unified network-level architecture supporting binocular convergence. However, we also observed slight asymmetries in the activity flow-generated activation strengths and other metrics explained throughout the study between the right and the left oculomotor processes, indicating that the contributions of each eye to binocular vision may not be fully symmetrical. While both eyes work in coordination to achieve a single binocular vision, the underlying distributed network dynamics may differ in magnitude or configuration for each eye. This insight is particularly relevant to dysfunctions of binocular eye movements. It raises important questions about whether the impairments affect network-level distributions of each eye equally, whether compensatory mechanisms exist to mitigate unilateral deficits, and at what point such compensation fails to maintain intact binocular vision. For example, the oculomotor dysfunction known as convergence insufficiency has shown that the peak speed of the left and right eye movement is asymmetrical and becomes more symmetrical with vision rehabilitation. Future research should investigate how distributed network mechanisms differ between the right and left eyes, whether they support eye-specific compensatory processes, and whether targeting the more affected eye’s network dynamics, rather than assuming symmetrical involvement, could provide more precise and effective clinical interventions, tailored for each participant’s needs.

While the involvement of visual networks in driving convergence-related activations within the CCC is expected, given the visual nature of the tasks used in our fMRI paradigm, the prominent role of the DAN holds particular significance for vision research. This finding highlights a potential bidirectional relationship between attentional systems and binocular eye movement control. Specifically, it may help explain why individuals with attention deficits, such as those with ADHD, often experience oculomotor deficits and difficulties with near-vision tasks like reading or using digital devices ^60^. Conversely, it also raises the possibility that individuals with binocular vision impairments may exhibit attentional disruptions due to impaired oculomotor coordination ^61^. In fact, studies indicate that patients diagnosed with ADHD are considerably more prone to being diagnosed with convergence insufficiency, and vice versa ^62^. These insights underscore the importance of considering attention-vision interactions in both clinical assessment and intervention strategies.

It is crucial to emphasize that the concept that vergence specificity and functional activations within the CCC are regulated by distributed large-scale networks does not diminish the significance of localized activations in oculomotor regions. In fact, this finding corroborates the notion that these localized activations are highly specific to the oculomotor regions, which are propagated through large-scale networks. This framework helps to explain how these local activations emerge in the brain regions associated with convergent eye movements. Therefore, this framework avoids perceiving local activations as mere local computations while there is clear evidence of a distributed process. Furthermore, the specificity of these regions when they receive information from propagated activations through large-scale networks shows their importance in winning the competition in performing local computations compared to other brain regions that receive the same propagation, probably by using inhibition. When a person performs a convergence task, the specificity of the oculomotor regions in receiving and processing information makes them unique in their computation of local activations, thanks to their distinct connectivity throughout the brain.

Finally, some alternative modeling approaches could be applied to the analytical framework used in this study. However, these extensions fall beyond the scope of the current manuscript and may detract from the central focus of our findings. First, task-specific FC could be estimated separately for each task condition to be used in the activity flow mapping. However, based on strong evidence that resting-state captures intrinsic brain organization that generalizes across a broad range of cognitive and perceptual states, we chose to use resting-state FC. Prior work has shown that resting-state FC closely resembles functional connectivity observed during task performance ^28^ and that resting-state networks spatially align with regions activated during task engagement ^13^. This allowed us to construct a unified, state-general model that avoids the need for computing separate connectivity patterns for each task condition and is not tied to any single stimulus or task context. Therefore, this emphasizes resting-state FC as a stable, reproducible feature, particularly valuable in longitudinal designs. To estimate these functional connections, we employed graphical lasso ^51^, a regularized form of partial correlation that addresses the limitations of both pairwise correlation and unregularized partial correlation ^52,63^. Unlike pairwise correlation, which cannot distinguish direct from indirect dependencies, FC calculated with regularized partial correlation using graphical lasso effectively isolates direct connections by suppressing spurious covariation. Moreover, compared to unregularized partial correlation, graphical lasso offers substantially improved test-retest reliability, robustness to noise and motion, and more accurate alignment with structural ground truth. Prior work has shown that this method provides stable, biologically meaningful network structures that predict task-evoked responses and behavioral variability ^51^. Therefore, our use of a graphical lasso to estimate resting-state FC yielded a stable and generalizable model of cortical communication that reliably supports distributed activity flow across task contexts, including those involving voluntary convergent eye movements. Second, although using resting-state FC provides extensive benefits in modeling task-evoked activations, it lacks directional information. While it reflects statistical associations between regions, it does not indicate the direction of influence. This could raise a potential causal confound in generative models, where observed activation patterns could, in theory, result from feedback signals originating in the target region itself rather than feedforward propagation from distributed sources. However, the design of the activity flow mapping framework helps to overcome this concern. Specifically, the target region is always held out of the activity flow model—that is, its own activation is not used to compute its generated activation. Instead, we rely solely on the activations of all other regions and their resting-state FC with the target. This holdout strategy ensures that feedback from the target region cannot influence the activity flow-generated activations, thereby reducing the risk of circularity or reverse causality in our inferences. As a result, our findings provide a more confident reflection of the feedforward contributions of distributed network interactions to the emergence of task-evoked activation patterns. Future work should investigate the causal direction and temporal dynamics of network interactions to better understand how distributed systems contribute to—or are influenced by—task-evoked activations in the CCC. Third, in the present study, we first defined the regions of interest as the CCC and then applied activity flow mapping within this localized circuitry. Future work could expand upon this approach by first identifying all cortical regions exhibiting task-evoked activation, potentially including areas outside the predefined circuitry, and applying activity flow mapping across these regions. This would allow for a more comprehensive assessment of both core convergence-related regions and additional contributing areas. Specifically, such an approach would benefit from larger sample sizes and could be further strengthened by incorporating individualized network parcellations ^64^ to account for inter-participant variability in functional architecture. This approach may also provide more precise correlations with clinical measures in populations with impaired binocular vision, better reveal the neural differences underlying functional impairments, and improve discrimination between clinical and healthy groups. Further, it could potentially delineate differences of convergence dysfunctions from abnormal development, brain injury such as concussion, or neurodegenerative diseases such as Parkinson’s Disease. Fourth, while our study successfully generated convergence-related activations with high spatial similarity to actual activation patterns, the magnitude of these generated responses was lower than that reported in prior activity flow mapping studies using large-scale cognitive datasets ^38^, but more comparable to results from clinical studies ^65^. This difference may be attributable to the high data quality and population-level diversity available in gold-standard datasets such as the Human Connectome Project. In contrast, our study focused exclusively on healthy individuals drawn from a controlled sample within a randomized controlled trial, lacking the broad inter-individual variability typically seen in population datasets. To mitigate potential confounds, we applied rigorous preprocessing and denoising procedures, allowing us to isolate the contribution of distributed functional connectivity to activation generation within the convergence system. Nonetheless, our use of a linear activity flow framework may have limited our ability to capture more complex or nonlinear representational dynamics, particularly in early visual regions or cross-network interactions. Incorporating nonlinear terms or adopting more expressive models that account for context-dependent modulation of resting-state connectivity may further improve the fidelity of activation generation in future work. This is especially relevant as we move toward clinical applications, where network topologies may deviate more substantially from healthy baselines.

This study has some limitations. First, our analysis was restricted to cortical regions, excluding subcortical and cerebellar structures that are known to contribute to oculomotor control and visuomotor integration. Future studies should include these regions to develop a more comprehensive understanding of the neural architecture supporting convergence and specifically explain the role of cerebellar regions in emerging convergent eye movements. Second, while our study focused on voluntary convergent eye movements, these movements differ significantly from involuntary, natural convergence behaviors encountered in everyday visual tasks. We encourage the development of innovative scanning protocols to address this in future studies. Third, our use of a single field map and relatively short resting-state acquisition (150 timepoints with a 2-second TR) could be enhanced in updated protocols to improve the reliability of FC estimates, particularly for fine-grained network modeling. Fourth, our sample size, though moderate, represents the upper limit of high-quality, single-site fMRI datasets with task and resting-state data collected under uniform protocols. Finally, the present study did not include cognitive task conditions beyond visuomotor and motor learning paradigms. While this design allowed us to isolate the network mechanisms specific to convergence eye movements, incorporating cognitive tasks in future work could provide a more comprehensive understanding of how the CCC interacts with broader cognitive control systems. This would be especially valuable when extending the framework to clinical populations, such as those with ADHD, post-concussion symptoms, or convergence insufficiency, where impairments may involve both oculomotor and higher-order cognitive dysfunction.

In summary, cortical activations during binocular convergent eye movements are shaped by distributed functional interactions across the visual and attention networks rather than arising solely from local activation within eye movement regions. We observed consistent bilateral engagement of the CCC, although with asymmetries suggesting that one hemisphere (or eye-specific network) may have a stronger influence. Among all networks, VIS1, VIS2, and the DAN contributed most prominently to convergence-related activation patterns, both during tasks specifically designed to evoke convergence and during unrelated visual and motor learning conditions. These findings support a model in which intrinsic resting-state FC organizes convergence control within a broader, domain-general network architecture. Future work should extend the methodology used here to clinical populations to evaluate the translational possibility of the distributed network mechanisms underlying convergent eye movements. This could show whether alterations in distributed and intrinsic resting-state FC underlie deficits in convergence control and whether these patterns differ systematically across diagnostic groups. Additionally, the framework can benefit longitudinal studies that evaluate therapeutic interventions. Furthermore, given the observed asymmetries between right and left eye processes in our results, future investigations should also explore whether one hemisphere or eye-specific network plays a dominant role in convergence-related activation and whether such asymmetries could serve as biomarkers for dysfunction or compensation.

## Methods

### Participants

This study is part of a randomized controlled trial, the convergence insufficiency neuro-mechanism in adult population study (CINAPS) ^66^. The study protocol followed the Declaration of Helsinki and was reviewed and approved by the Institutional Review Boards of the New Jersey Institute of Technology and Rutgers University (<u>ClinicalTrials.gov</u> registration Identifier: NCT03593031). All participants gave signed, written informed consent. The prior protocol paper provided detailed information about the study design, methodology, and participant selection for the project ^66^. In brief, the study population consisted of participants with binocularly normal vision, visual acuity of 20/25 or better with refractive correction, local stereopsis of 70 seconds of arc, and global stereopsis of 250 seconds of arc. Individuals with a history of head injury or concussion and those with any neurological or retinal disease were excluded. Data collection was performed at the Rutgers University Brain Imaging Center on 50 participants. Technical errors in saving the file structure of the imaging datasets in one of the tasks resulted in the removal of two participants from further analysis. The dataset used in the following study comprises resting-state and task-evoked fMRI scans from 48 healthy young participants with binocularly normal vision. The average age was 21.8, and participants were between 18 and 34 years (15 female). Of these participants, 27 had additional repeat data from the resting-state scans, with a time interval of 1– 31 days (mean 7.67 ± 7.72 days) between the two scans, depending on the participant’s and imaging center’s availability. The repeat scans were typically scheduled about two weeks after the initial scan.

### Task Paradigms

Convergent eye movements were studied using the vergence motor task (VM). While the main task of interest in the present study was VM, other tasks used in the following analysis (in particular, the response profile in the network contribution analysis) were utilized as control tasks to the VM task. Due to the primary focus on convergent eye movements and allowing participants to rest their eyes while performing them, two runs of the VM task were collected. All participants received training on the tasks before the fMRI session to ensure they could perform the tasks effectively in the scanner. The details of each task are described below and are adapted from previous studies ^48,66^.

#### Vergence Motor Task

The VM task included symmetrical convergent eye movements embedded in the alternating blocks of convergent eye movements, followed by rest periods. Task blocks included visual stimulations for performing VM tasks with high-frequency repetitions (VM-many, eight occurrences of convergence movements) and low-frequency repetitions (VM-few, four occurrences of convergence movements). These two blocks were initially designed to investigate the effective number of VM movements to exert reliable fMRI activations ^48^. However, for the current study, both VM-many and VM-few task blocks served the same purpose of studying convergent eye movements. In each run of the VM task, there were five blocks of VM-many and five blocks of VM-few. The VM-many blocks stimulated eight vergence eye movements and lasted for 19 seconds each, while the VM-few blocks stimulated four vergence eye movements and lasted for 18 seconds each. After preprocessing, the denoised data from the two VM runs were concatenated (described below).

#### Vergence Sensory Task

The Vergence Sensory (VS) task was designed to provide sensory stimulation. Participants were instructed not to perform convergent eye movements while viewing the visual stimuli associated with the convergent eye movement tasks. This task was initially intended to study the differences between the afferent and efferent pathways of convergent eye movements in comparison to the VM task. Similar to the VM task, there were two types of task blocks for the VS task: VS-many with eight occurrences of stimuli and VS-few with four occurrences of stimuli. Besides the initial intention of the VS task design for the current study, both VS-many and VS-few task blocks served the same purpose as the control task. In each VS task, there were five blocks of VS-many and five blocks of VS-few. The VS-many blocks lasted for 22 seconds each, while the VS-few blocks lasted for 18 seconds each.

#### Saccadic Motor Task

The Saccadic Motor (SM) task was designed to study the saccadic eye movements in the original study. Participants were instructed to follow a square moving to the right and left on the screen to capture their ability to perform saccades. There were two types of task blocks for the SM task: SM-fast with 24 occurrences of saccadic stimuli and SM-slow with 12 occurrences of saccadic stimuli. Similar to the VM and VS tasks, both SM-many and SM-few task blocks served the same purpose as the control task for the current study. There were five blocks of each task type in each SM task, and both lasted 24 seconds.

#### Finger Tapping

The Finger Tapping (FT) task was implemented as a control measure to assess participants’ overall motor responses. Specifically, the FT task acted as a control for assessing motor performance, excluding the oculomotor system. Participants were instructed to perform finger tapping at both slow and fast speeds, as guided by the instructions displayed on the screen. Therefore, there were two types of task blocks for the FT task: FT-fast and FT-slow, with each block occurring three times and lasting 20 seconds. Similar to the VM, VS, and SM tasks, the FT-fast and FT-slow blocks were designed to serve as a control task for this study.

### Data Acquisition

Data Acquisition was conducted at a 3T Siemens TRIO (Siemens Medical Solutions, USA) with a 12-channel head coil. Before each scan, the experimental protocol, the scanning process, and the safety precautions were explained to the participants. Each participant was instructed to lie still and was reminded not to move their head during the scan while staying awake. All participants were asked to remove any metal or ferrous materials before entering the scanning room. Additionally, spongy pads were placed around their head inside the head coil to help minimize head movement.

A single echo planar imaging (EPI) sequence was used to obtain functional images with repetition time (TR) = 2000 ms, echo time (TE) = 13 ms, field of view (FOV) = 192 mm, flip angle = 60°, total number of acquired axial slices = 53, and 3 mm isotropic voxels. A high-resolution T1-weighted anatomical scan was also collected with 1 mm isotropic voxels using a magnetization-prepared rapid acquisition gradient-echo (MPRAGE) sequence. Participants closed their eyes during the anatomical scan. A spin-echo fieldmap was collected in the posterior-to-anterior direction. The resting-state scan was collected for 5 minutes (150 TRs). VM task run-1 and run-2, VS and SM scans were collected for 6 minutes and 56 seconds (208 TRs). FT scan was collected for 3 minutes and 4 seconds (92 runs). Participants completed all tasks in a single fMRI session, with each task scan collected consecutively and separated by brief breaks.

#### fMRI Preprocessing

Before preprocessing, we checked all structural and functional data for any apparent issues or unusual patterns. Preprocessing of the anatomical and functional images was performed using fMRIPrep 24.0.1^67^, which is based on Nipype 1.8.6 ^68^. Details of the fMRIPrep pipeline are given in the supplementary information.

Following minimal preprocessing with fMRIPrep, we continued further analysis with surface data in CIFTI format. Each functional image was projected onto the fsLR cortical surface grayordinates files ^69^, containing 91k samples (32k vertices per hemisphere). The advantages of using grayordinates and CIFTI format have been extensively explained in prior research ^70,71^. We then downsampled the graordinate data into the Glasser surface-based cortical multimodal parcellation (MMP) atlas ^49^. This parcellation integrates various neuroimaging modalities to enhance the accuracy of assigning specific regions of the cortex. As a result, it offers a systematic approach to dividing the cortex into a manageable number of functionally relevant units, which helps reduce the number of statistical comparisons. Next, further standard preprocessing steps were applied to the parcellated task and resting-state data. We demeaned and detrended the task and resting-state data and performed nuisance regression. For resting-state scans, we removed the first five TRs. The nuisance regression followed previous studies based on Ciric et al ^20,72^. We performed nuisance regression to remove motion and physiological artifacts. Six motion parameters, their derivatives, and quadratics (24 total) were modeled for motion artefacts. Anatomical CompCor (aCompCor; ^73^) within fMRIprep was utilized on white matter and ventricle time series to model physiological noise. Signals of the five principal components, along with their derivatives and the quadratics, were modeled and removed, totaling 20 physiological noise regressor parameters. Additionally, for the resting-state data, we also calculated relative root mean square displacement to identify high movement frames in the data (with >0.25 mm frame-wise displacement) and added an additional “spike” regressor to the nuisance regression. We employed nuisance regression using aCompCor, which includes components that are similar to the global signal but has the advantage of not removing gray matter signals ^20^. Consequently, we chose not to eliminate the global signal, as there is substantial evidence indicating that its removal introduces negative correlations into the data and not a standard practice for task activation analyses ^74,75^.

### fMRI task activation estimation

Cortical activations associated with task-state fMRI conditions were estimated using a standard general linear model (GLM). This model convolved the timing of tasks (for each block-design condition) with the canonical hemodynamic response function from SPM ^76^. The result was a matrix of regression coefficients with dimensions of 360 regions by conditions for each task, representing the activation amplitudes for each participant.

### Resting-state functional Connectivity Estimation

Functional connectivity (FC) was estimated using regularized partial correlation, implemented via graphical lasso, to characterize brain region interactions reliably while mitigating confounding influences. Specifically, each participant’s preprocessed resting-state fMRI data were analyzed individually, where graphical lasso was applied with cross-validation to optimally select the L1 regularization parameter. This cross-validation-based regularization effectively reduced overfitting to noise, thus enhancing the reliability and validity of FC estimates compared to standard pairwise and unregularized partial correlations. Empirical resting-state time series data for each participant were input to the graphical lasso algorithm, resulting in participant-specific FC matrices capturing robust and direct functional connections between pairs of brain regions. Prior research ^51^ has demonstrated that graphical lasso provides superior reliability, robustness to common fMRI artifacts such as participant motion, and enhanced correspondence with structural connectivity and task-evoked activations. Consequently, graphical lasso-derived FC matrices were used as inputs for subsequent network analyses, including activity flow mapping, to ensure a more valid estimation of regional interactions.

### Functional activations generated via activity flow

We employed activity flow mapping to demonstrate that interactions across the entire cortex can accurately represent localized, convergence-specific functional responses within the Cortical Circuitry of Convergence (CCC). This modeling approach maps brain activity by leveraging the inherent relationship between activation patterns and FC. Originally developed by Cole et al. ^35^, this method has been extensively utilized and described in previous research ^38^, including a protocol paper ^37^. The approach utilizes a specific formula to simulate brain activity in a target region based on the activity flows from other regions and their connections to the target area:

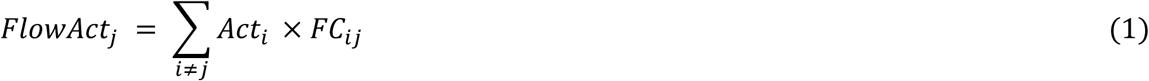

Where Act_i_ is the functional activity of the source region i (β values estimated by a GLM); FC_ij_ is the functional connectivity (here with graphic lasso) between the target brain region j and all other brain regions (i). FlowActⱼ is the activity flow-generated functional activity of the target region/parcel j. Here, j is the target brain region (held out), and i is all other brain regions of the parcellation.

Using this formula in any given task allows for mapping and generating functional activity in a target region by summing the actual functional activity of all other source brain regions weighted by the estimated FC between the target and source regions. In summary, activity flow mapping proposes that task-evoked functional activity in a given task is propagated between brain regions via their FC. The activity flow mapping procedure tests the hypothesis that the total activity in source regions, based on brain connectivity patterns, matches that of target regions. It has been found that this mapping is most accurate with high global coupling and low self-coupling in simulated models ^35^. This indicates that successful activity flow mapping in empirical brain data likely retains these high global coupling properties, suggesting that it effectively reflects the contribution of distributed network interactions to specific activations. To avoid circularity in generating activations that could be influenced by the regions responsible in convergent eye movement (explained in the next section), we excluded all these regions from the source set when estimating activation for each target region. This ensured that activity flow-generated activations were not driven by direct contributions from other functionally related regions involved in convergent eye movements.

To evaluate the accuracy between actual empirical activations and activity flow-generated functional activities, these values were compared using the Pearson correlation coefficient (Pearson r), the coefficient of determination (R²), and mean absolute error (MAE). Each of these comparison metrics has distinct advantages and disadvantages, but together, they offer a comprehensive characterization of mapping accuracy. These comparison statistics were calculated for each participant and then averaged across participants to provide a random effects estimate (see the protocol paper for more details ^37^).

We further refined this accuracy evaluation to the convergence motor task (VM) to compare activity flow-generated versus actual activation patterns across all cortical parcels during the VM task conditions across participants.

### Cortical Circuitry of Convergence

Previous research has shown that distributed network interactions can accurately map localized, convergence-specific functional responses ^35,77,78^. To testify this in our dataset, we centered our analysis on the functional brain regions previously identified by Morales et al. ^48^, prioritizing regions exhibiting high intraclass correlation for reliably capturing activity associated with convergent eye movements. Typically, direct registration of individual participants to a standard coordinate system is preferred. However, as previously published results utilized the Montreal Neurological Institute (MNI) space, we employed the Glasser atlas in MNI space to facilitate alignment with the convergence-specific ROIs ^79^. We then used MNI coordinates corresponding to each ROI and mapped these onto the nearest matching regions within the volumetric Glasser atlas. Consequently, we identified the closest Glasser MMP parcels for each ROI coordinate set. Additionally, we integrated the Cole-Anticevic brain-wide network partition (CAB-NP) ^50^ for network assignments. The CAB-NP partition, derived from Human Connectome Project resting-state fMRI data, employs a Louvain community detection algorithm to classify MMP parcels into 12 distinct functional networks. Table 1 shows Brain Regions in the CCC, their corresponding indices in the MMP atlas, network assignment in the CAB-NP, and MNI coordinates in the volumetric space.

**Table 1.** Brain Regions in the Cortical Circuitry of Convergence.

| MMP Atlas Region name | MMP Atlas indices (L, R) | Network assignment | Left Hemisphere MNI Coordinates | Right Hemisphere MNI Coordinates |
| --- | --- | --- | --- | --- |
| Primary Visual Cortex | 1, 181 | Visual-1 | -10 -88 6 | 10 -88 6 |
| Frontal Eye Field | 10, 190 | Cingulo Opercular | -28 -6 48 | 28 -6 52 |
| Supplementary and Cingulate Eye Field | 43, 223 | Cingulo Opercular | -6 8 50 | 6 8 50 |
| Anterior IntraParietal Area | 117, 297 | Dorsal Attention | -30 -48 50 | 30 -48 50 |

After defining regions within the CCC, we extended the whole-cortex accuracy evaluation and tested our model more directly on the brain regions involved in convergent eye movements. We refined our analysis, focused on the CCC across all eight task conditions. These conditions were used to probe both convergence-related processes (including the VM and VS tasks) and non-convergence-related functions (including SM and FT). Therefore, we quantified a response profile of the CCC, also known as population receptive field analysis ^80^. This response profile was defined as the task-evoked activations of the CCC across all eight task conditions. This profile enabled us to verify convergence-specific responsiveness within the CCC and demonstrate that activity flow mapping reliably captured activation patterns across both convergence-related and non-convergence-related motor tasks. This supports the notion that activity flow mapping in these regions generalizes across a broad range of visual and non-visual motor demands.

To further test whether the CCC is selectively responsive to the motor execution of convergence rather than passive visual input, we compared activation levels during the VM with those from a matched sensory control task (VS), in which participants viewed the same stimuli without performing convergence. This analysis was performed on both actual and activity flow-generated activations. If these regions are specifically involved in generating convergence eye movements, we expected greater activation during VM relative to VS, reflecting their sensitivity to the motor, rather than purely visual, demands.

To avoid circularity in activity flow-generated activation that the CCC itself could influence, we excluded all other CCC regions from the source set when estimating activation for each target region. This ensured that mapping was not driven by direct contributions from other functionally related regions involved in convergence eye movements.

#### Vergence Specificity

To determine how specifically regions within the CCC capture convergent eye movements, we quantified the vergence specificity index as the ratio of mean activation amplitude during the VM task conditions relative to the non-vergence sensory task conditions (i.e., VS). The VS task was selected as an ideal control because it explicitly required the absence of convergence, thus isolating convergence-related activity in the VM condition. Using a ratio rather than a difference score minimizes sensitivity to variations in β value scales across brain regions.

Prior to computing this ratio, we normalized activation values using a standard min-max normalization (feature scaling) to ensure interpretability and comparability across participants by eliminating negative values and rescaling all activation values to a common [0, 1] range. This normalization was performed per participant based on their minimum and maximum activation values across all eight task conditions within the CCC. Vergence specificity was then computed separately for activity flow-generated and actual data per participant and averaged across regions within the CCC.

We computed vergence specificity with the following formula:

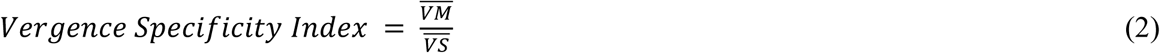

where 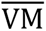 denotes the mean activation amplitude during convergence motor responses and 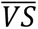 denotes the mean activation amplitude during non-convergence sensory responses.

A vergence specificity score of 1.0 reflected the null hypothesis, indicating equal responses to VM and VS conditions. Scores greater than 1.0 (e.g., *n*) showed that activation amplitudes during convergence tasks were *n* times greater than during control sensory task conditions. For computing vergence specificity, we identified and removed outlier participants using the median absolute deviation method with a conservative ±5 deviation threshold, as described by Leys et al.^81^.

Since normality tests indicated that the data deviated from a normal distribution in some of the variables (p < 0.05), we conducted non-parametric permutation testing to ensure robust inference, as recommended in neuroimaging studies, due to its minimal assumptions about data distribution and its superior control over false positives. Specifically, we employed upper-tailed one-sample t-tests using nonparametric permutation-based significance testing with a maximum statistic (max-T) correction to account for multiple comparisons, using 100,000 permutations to enable p-value estimation as low as 0.00001, in accordance with established practices in fMRI analysis ^82,83^. This method provides a conservative and assumption-free estimate of statistical significance. We used the max-T nonparametric test here to evaluate the statistical significance of the difference in vergence specificity against the null hypothesis of 1.

#### Whole cortex contribution to the vergence specificity

To evaluate the extent to which whole-cortex distributed network interactions contribute to convergence-specific responses, we computed the percentage of actual vergence specificity (representing both distributed and localized processes) explained by activity flow-generated activity vergence specificity (representing distributed processes only). The remaining unexplained portion likely reflects local processing or other model-related errors, such as noise. If the mapped percentage exceeded 50% significantly, we inferred that vergence specificity in the CCC was primarily supported by distributed activity flow processes, recognizing this as a conservative (lower-bound) estimate. We used the max-T nonparametric test with 100,0000 permutations to evaluate the statistical significance of the difference in vergence specificity against the null hypothesis of 50%.

### Network Analysis

#### Network-level functional activations

We computed the role of each functional network in generating functional activities within the CCC. To test and compute this, we use the values that were employed to compute the activity flow-generated functional activity in each brain region. Since each functional activity was generated by the weighted FC of that region with the functional activity of all other regions in the brain, the amount of this flow of activity could be attributed based on the classification of each brain region into one of the functional networks defined in the CAB-NP. Therefore, averaging all flows at the network-level before reaching the held-out regions could provide the overall contribution of that network to the activity flow generation for the functional activity in the held-out region. Moreover, this process can be repeated for each region in the CCC to reach the network-level generated activities within the circuitry. We took the average of the flows for the regions inside the CCC. Finally, we conducted a statistical comparison of these network contributions to identify a hierarchical contribution of networks to the activity flow mapping towards the CCC. This comparison provided insights into which network outperformed others in generating functional activity within the CCC. We used Max-T with 100,0000 permutations to address multiple comparisons in the network-level analysis.

Next, we computed the contribution of each functional network to the response profile of the CCC. This means calculating the activity flow mapping of the CCC for all task conditions restricted by the CAB-NP functional networks. Consequently, we obtained values for each network that summed to the whole-model activity flow mapping. These values represented the activity flow-generated activations broken down by CAB-NP networks in the source regions, and this approach effectively parsed the activity flows into their network components. We further testify that the sum of all these network-level values equals the original full model values. Next, to compute the network-level contributions to the activity flow-generated activities across the CCC in the response profile, we conducted a dominance analysis ^84^. Dominance analysis is used to determine the relative importance of predictors for both regression and classification models. Here, the “relative importance” is based on the additional contribution of a predictor in all subset models. The purpose of determining predictor importance in the context of dominance analysis is not to select a model but rather to uncover the individual contributions of the predictors. Dominance analysis in our study determines the dominance of one predictor (network-level activity flow predictions) over another by comparing their incremental explained variance (R²) contribution across all subset models. This method provides a robust, multicollinearity-resistant measure ^85^ of how different functional networks shape the activation profile of convergence-related regions through a distributed activity flow. Then, we performed multiple linear regression for each participant across conditions, using 12 network-level genrated activations as predictor variables and the actual activations as the response variable. The full-model R² represents the extent of variance in the response profile calculated from the actual functional activity that is explained by the response profile calculated from the network-level, activity flow-generated activations. After obtaining the full model R², we continued with subset models in the dominance analysis. This means a subset of multiple regressions with all possible combinations of predictors (network-based activity flow-generated activations). Since the whole model included 12 network-level predictors, this combination ranged from 1 to 11. Using a standard combination calculation, nCr, 4095 subsets of models were possible, where n was 12 and r was iterated between 1 and 11. This analysis provided the incremental contribution of each predictor to R² (network-based activity flow-generated activations). Subsequently, the average of these incremental contributions across models was considered as the partial R2 for each network, with the cumulative total equating to the R2 of the full model.

Furthermore, we assessed group-level differences in flows to the left and right CCC for VM and VS conditions using the actflow product (task activation × resting-state FC) at both the network and parcel levels. Activity flows were computed separately for VM and VS task conditions using the dot product of task-evoked activation and resting-state FC (i.e., the actflow term). To assess condition differences, we averaged flows into the CCC for each subject, then performed a paired, upper-tailed max-T permutation test (100,000 permutations) across the 12 CAB-NP networks. For parcel-wise comparisons, we extracted subject-wise flow values from each of the 360 Glasser parcels into the CCC and computed paired differences between VM and VS. A separate max-T test was then applied across parcels to identify statistically significant effects while controlling for multiple comparisons.

### Repeated Resting-State Scan

We applied the full set of analyses described above to a subsample dataset from 27 participants. Following identical preprocessing steps as in the primary analysis, we replaced the resting-state functional data from each participant’s repeat scan and used it to repeat the generation of task-evoked activations. That is, for each participant, resting-state FC from the second scan was used to model functional activations observed in the original data. This approach aligns with our hypothesis that activity within the CCC can be generated from distributed activity in other brain regions, weighted by their resting-state FC to the CCC. Given that resting-state FC is typically stable across sessions in healthy individuals ^57^, this repeat analysis served to further validate our hypothesis regarding the contribution of distributed networks to convergence-related functional responses shaped by the resting-state FC signatures. In addition, we extended our analysis to include vergence specificity, network contributions to the CCC, and dominance analysis in both hemispheres.

### Control Analyses for Circularity Concerns and Potential Inferential Limitations

We applied some steps to address potential circularity in generating activation that could be influenced by the regions within the CCC. The first and critical step in this regard was excluding all these regions from the source set when estimating activation for each target region. This ensured that activity flow-generated activations were not driven by direct contributions from other functionally related regions involved in convergent eye movements.

#### Spatial Circularity Control in Activity Flow Mapping

fMRI data are known to demonstrate spatial smoothness in the range of 2–5 mm, largely driven by vascular hemodynamics rather than neural sources ^65,86^. This smoothness may potentially introduce bias into the generation of activity flow by permitting signals from the target region to leak into neighboring source regions, thereby violating the independence assumption inherent in generative modeling. To minimize this risk of circularity, we implemented a control analysis as a spatial exclusion procedure in the FC estimates. Specifically, for activity flow generations in each to-be-computed target region, we excluded all cortical parcels with any surface vertices located within 10 mm distance on the brain surface and masked the connections between each target and its spatially adjacent parcels. This control analysis was in addition to excluding eight anatomically defined ROIs from participating as source region in any target within the CCC. This dual masking strategy ensured that both spatially contaminated and task-relevant co-targets were removed from the source set during model evaluation in the control analysis.

#### Circulating the signal with the CCC to improve the generated activations

We further explored additional steps to improve the activity flow-generated activations within the CCC. One such option involved modeling potential recurrent signal propagation within the CCC itself as proposed by ^58^ in capturing the dynamics of the human visual system. To implement this, we conducted a secondary round of activity flow mapping constrained to the CCC regions. For each region with the CCC, the initially activity flow-generated activations from the whole-cortex model served as inputs. Each CCC region was then held out in turn, and the activation was generated using the remaining seven CCC regions and their FC estimate weights. This step simulated a single round of intra-complex activity flow, allowing to examine whether reciprocal interactions within the CCC could refine the generation of task-evoked responses. The decision on model performance and improvement was evaluated by comparing secondary-generated activations to actual activations using *R²*, and vergence specificity measures. Improvements relative to the primary model were used to assess whether internal signal circulation added explanatory value. To achieve model comparison values, iterations were conducted over held-out regions, task conditions, and participants.

#### Condensed Cortical Models

To test whether a restricted set of functional networks could account for convergence-related activation in the CCC, we constructed and evaluated several submodels of the full whole-cortex activity flow framework. These models tested whether activity propagation limited to one or a few networks could replicate the model accuracy and vergence specificity achieved by the whole-cortex distributed model.

Each submodel included a specific combination of cortical networks defined by the CAB-NP atlas, with source regions limited to the selected networks and the CCC regions (when applicable). Five submodels were evaluated: (1) VIS-CON-DAN (Visual-1, Visual-2, Cingulo-Opercular, and Dorsal Attention), (2) VIS-CON, (3) VIS-DAN, (4) VIS-Only, and (5) DAN-Only. Additionally, we tested a sixth model, V1-Initiated, in which the only source regions were left and right V1. In all submodels (except V1-Initiated), all CCC region was held out, and their generated activations were computed using the remaining source regions’ task activations weighted by their resting-state FC estimates. For the V1-Initiated model, we first mean-centered the activation values of left and right V1 to eliminate any convergence-related signal potentially influenced by distributed activity or CCC feedback. These centered V1 activations were then used as source regions to generate activations in the remaining six CCC regions. Mean-centered activations in each V1 region were used to generate activations in the other hemisphere’s V1.

For each model, we assessed vergence specificity. Additionally, we quantified model fit by computing R² values between generated and actual response profiles within the CCC, separately for the left and right hemispheres. R² was calculated by iterating over held-out regions, task conditions, and participants. The performance of each submodel was compared to the primary whole-cortex model to evaluate whether restricted connectivity could sufficiently capture convergence-related activations.

## Data Availability

The data used in this study are a part of a randomized controlled clinical trial, and the public repositories are not publicly available currently. The IRB during data collection did not ask participants to post non-identifiable information publicly, hence data will be shared upon request.

### Code Availability

Code to conduct activity flow mapping and the subsequent statistics is publicly available via the Brain Activity Flow (“Actflow”) Toolbox (https://colelab.github.io/ActflowToolbox/). The code used in this study is leveraged from the code originally available via (https://colelab.github.io/ActFlowCategories) from Cocuzza et al. ^38^, and modified for the present study. Connectome workbench is publicly available via (https://www.humanconnectome.org/software/connectome-workbench). The Cole-Anticevic Brain-wide Network Partition (CAB-NP) is publicly available via (https://github.com/ColeLab/ColeAnticevicNetPartition).

## Notes

### Competing Interest Statement

The authors have declared no competing interest.

