## Supplementary Information for "Distributed Cortical Network Dynamics of Binocular Convergent Eye Movements in Humans"

### fMRIPrep Preprocessing Pipeline

Preprocessing of the anatomical and functional images was performed using fMRIPrep 24.0.1[1], which is based on Nipype 1.8.6 [2]. A B0 nonuniformity map (or fieldmap) was estimated from the phase-drift map(s) measure with two consecutive GRE (gradient-recalled echo) acquisitions. The corresponding phase-map(s) were phase-unwrapped with prelude (FSL 6.0.7.7).

#### Anatomical data preprocessing

From the original longitudinal study, a total of 2 T1-weighted (T1w) images were available for each participant and used for anatomical data preprocessing. Each T1w image was corrected for intensity non-uniformity (INU) with N4BiasFieldCorrection, distributed with ANTs 2.5.1 [3,4]. The T1w-reference was then skull-stripped with a Nipype implementation of the ANTS brain extraction workflow. An anatomical T1w-reference map was computed after registration of 2 T1w images (after INU-correction) using mri_robust_template and brain surfaces were reconstructed using FreeSurfer 7.3.2 [5,6], and the brain mask estimated previously was refined with a custom variation of the method to reconcile ANTs-derived and FreeSurfer-derived segmentations of the cortical gray-matter of Mindboggle [7]. Volume-based spatial normalization to two standard spaces (MNI152NLin2009cAsym, MNI152NLin6Asym) was performed through nonlinear registration with antsRegistration, using brain-extracted versions of both T1w reference and the T1w template. Grayordinate "dscalar" files containing 91k samples were resampled onto fsLR using the Connectome Workbench [8].

#### Functional data preprocessing

For each functional image, the following preprocessing was performed. First, a reference volume was generated using a custom methodology of fMRIPrep for use in head motion correction. Head-motion parameters with respect to the BOLD reference (transformation matrices, and six corresponding rotation and translation parameters) were estimated before any spatiotemporal filtering using mcflirt [9]. The estimated fieldmap was then aligned with rigid-registration to the target EPI (echo-planar imaging) reference run. The field coefficients were mapped onto the reference EPI using the transform. The BOLD reference was then co-registered to the T1w reference using bbregister (FreeSurfer), which implements boundary-based registration [10]. Co-registration was configured with six degrees of freedom. Several confounding time-series were calculated based on the preprocessed BOLD data: framewise displacement (FD), DVARS, and three region-wise global signals. FD was computed using two formulations following Power (absolute sum of relative motions, [11]) and Jenkinson (relative root mean square displacement between affines, [9]). FD and DVARS were calculated for each functional run, both using their implementations in Nipype (following the definitions by Power et al. 2014). The three global signals were extracted within the CSF, the WM, and the whole-brain masks. Additionally, a set of physiological regressors were extracted to allow for component-based noise correction

(CompCor, [12]). Principal components were estimated after high-pass filtering the preprocessed BOLD time-series (using a discrete cosine filter with 128s cut-off) for the two CompCor variants: temporal (tCompCor) and anatomical (aCompCor). tCompCor components were then calculated from the top 2% variable voxels within the brain mask. For aCompCor, three probabilistic masks (CSF, WM and combined CSF+WM) were generated in anatomical space. The implementation differs from that of Behzadi et al. in that instead of eroding the masks by 2 pixels on BOLD space, a mask of pixels that likely contain a volume fraction of GM is subtracted from the aCompCor masks. This mask is obtained by dilating a GM mask extracted from the FreeSurfer's aseg segmentation, and it ensures components are not extracted from voxels containing a minimal fraction of GM. Finally, these masks were resampled into BOLD space and binarized by thresholding at 0.99 (as in the original implementation). Components were also calculated separately within the WM and CSF masks. For each CompCor decomposition, the k components with the largest singular values are retained, such that the retained components' time series are sufficient to explain 50 percent of variance across the nuisance mask (CSF, WM, combined, or temporal). The remaining components were dropped from consideration. The head-motion estimates calculated in the correction step were also placed within the corresponding confounds file. The confound time series derived from head motion estimates and global signals were expanded with the inclusion of temporal derivatives and quadratic terms for each [13]. Additional nuisance timeseries were calculated by means of principal components analysis of the signal found within a thin band (crown) of voxels around the edge of the brain, as proposed by [14]. The BOLD time-series were resampled onto the left/right-symmetric template "fsLR" using the Connectome Workbench [8]. To project BOLD timeseries onto the fsLR cortical surface, grayordinates files [8] containing 91k samples (32k vertices per hemisphere) were generated with surface data transformed directly to fsLR space.

**Figure S1. FC estimates. A:** Single-participant FC estimate with regularized partial correlation. **B.** Single-participant FC estimate with Pearson correlation. **C.** Group-average FC estimates with Pearson correlation.

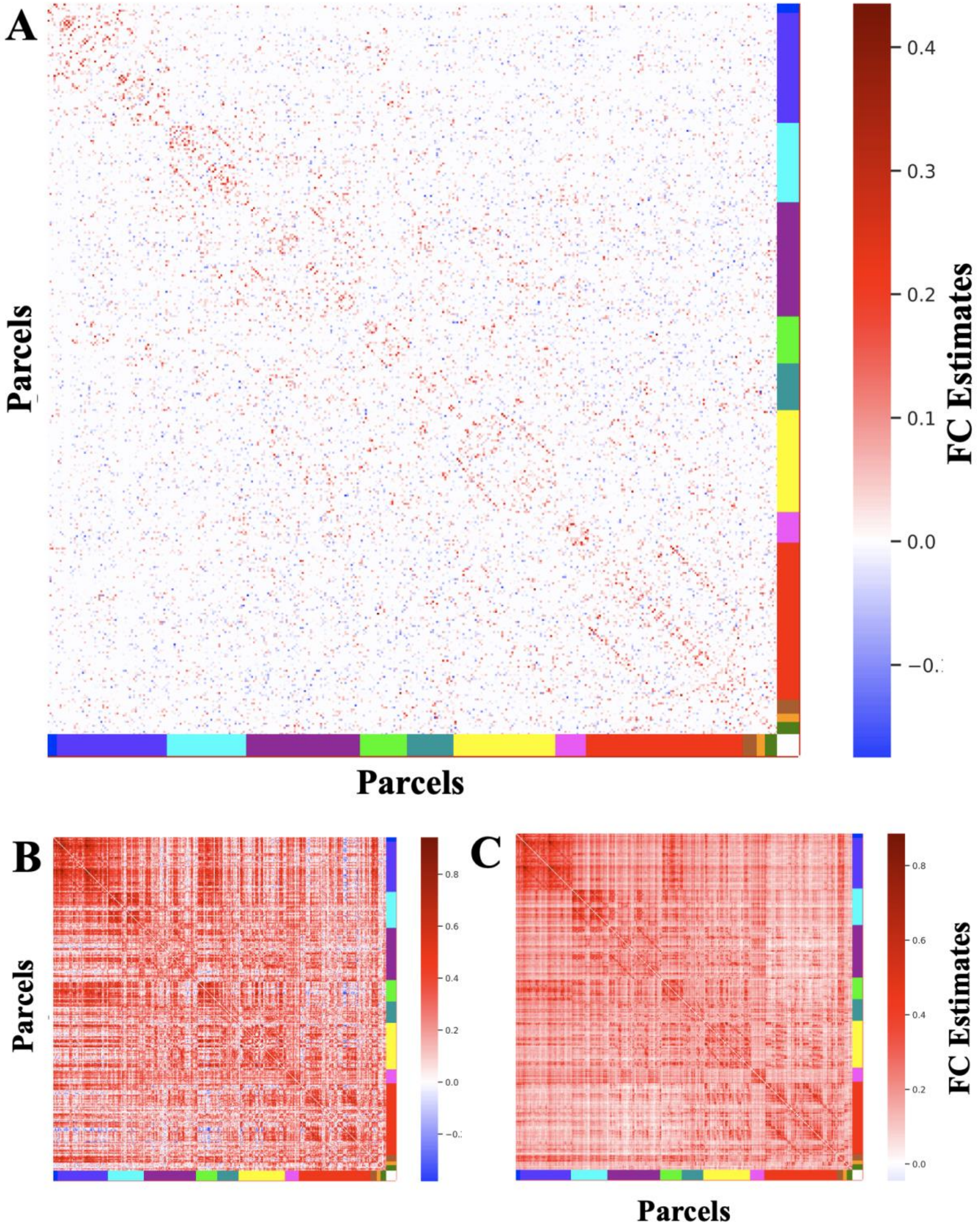

**Figure S2. Visualization of the overall model accuracy.** Comparison between actual and activity flow-generated activations across all conditions and brain parcels (360 cortical parcels from the Glasser MMP atlas, assigned to 12 networks by CAB-NP). Model accuracy was quantified as the similarity between actual and activity flow-generated activation patterns, and results were averaged across participants to assess consistency at the group level. This yielded a high overall model accuracy, as indicated by a Pearson's r of 0.75, an $R^2$ of 0.55, and an MAE of 0.99.

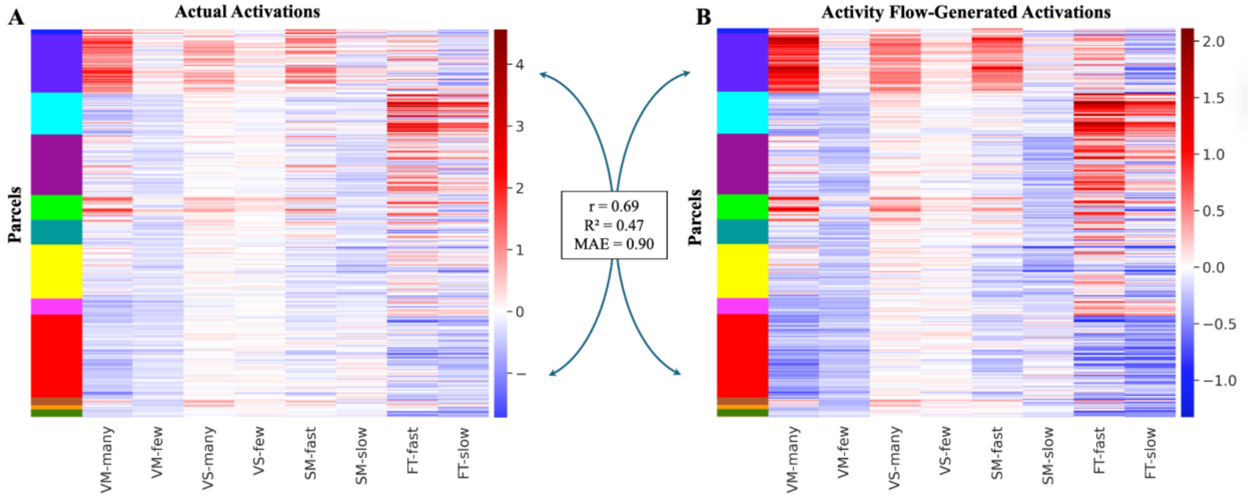

**Figure S3. Actual and activity flow-generated activation maps for all task conditions.** Actual (left panel) and activity flow-generated activations (right panel) are shown separately for each task condition. Comparisons in each row show the condition-wise similarity between the actual and activity flow-generated activations. VM: Vergence Motor task; VS: Vergence Sensory Task; SM: Saccadic Motor Task; FT: Finger Tapping Task (See Methods for details).

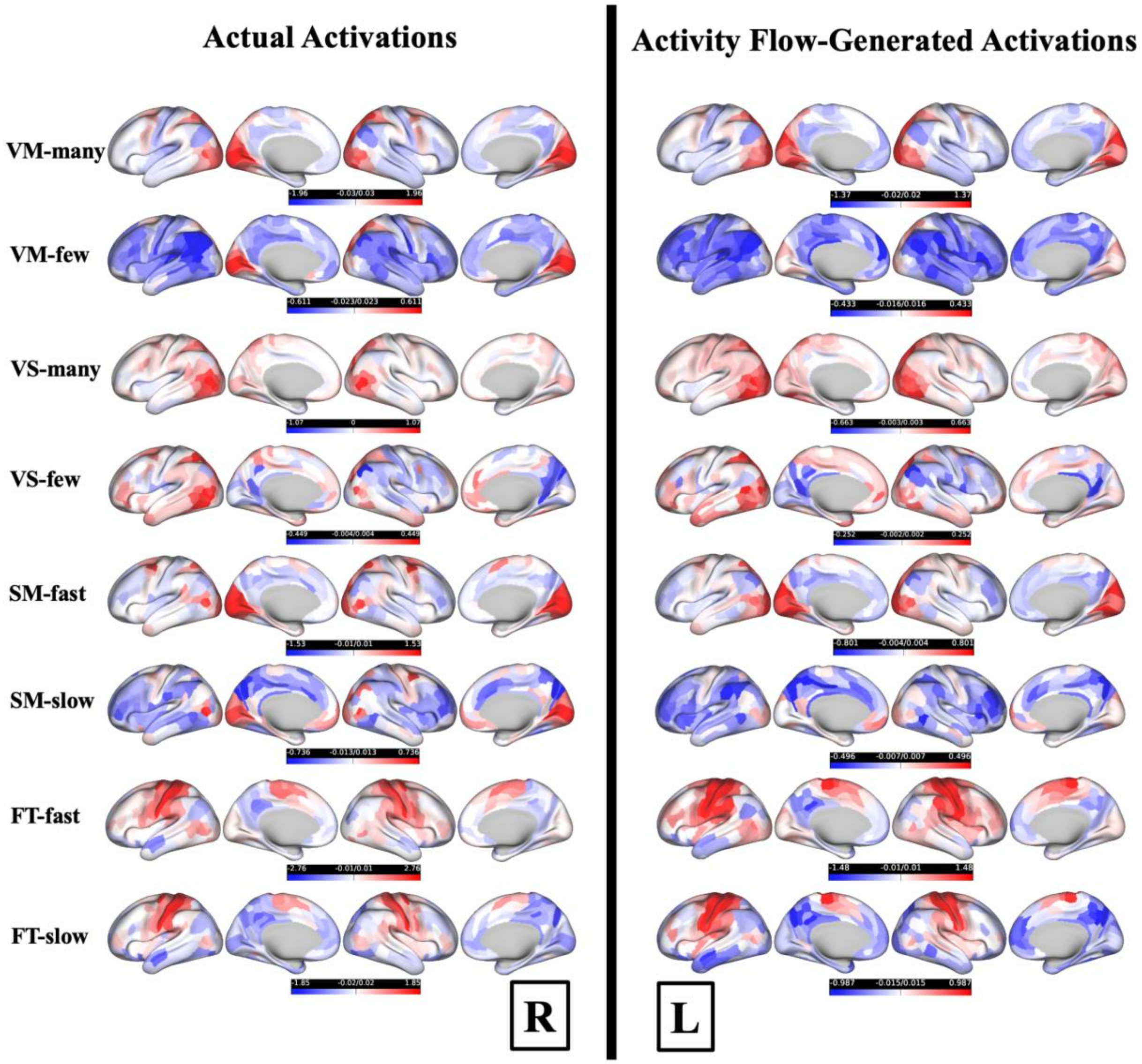

**Figure S4. Full matrix of observed t-statistics for pairwise network comparisons for the left and right hemispheres, in comparisons of the contributions of large-scale functional networks.** Each cell reports the t-statistic for the comparison between the row network and the column network (Row minus Column). Positive t-values (shown in shades of red) indicate greater activation in the row network relative to the column network. Negative t-values (shown in shades of blue) indicate greater activation in the column network relative to the row network. The diagonal is masked as it represents self-comparisons. Significance was determined based on a max-T corrected threshold (t = 3.4; p < 0.00001, 100,000 permutations). Significant comparisons, defined as t-values exceeding the threshold, are indicated with an asterisk (*). Directionality matters: for example, while VIS2→CON is significant, CON→VIS2 is not.

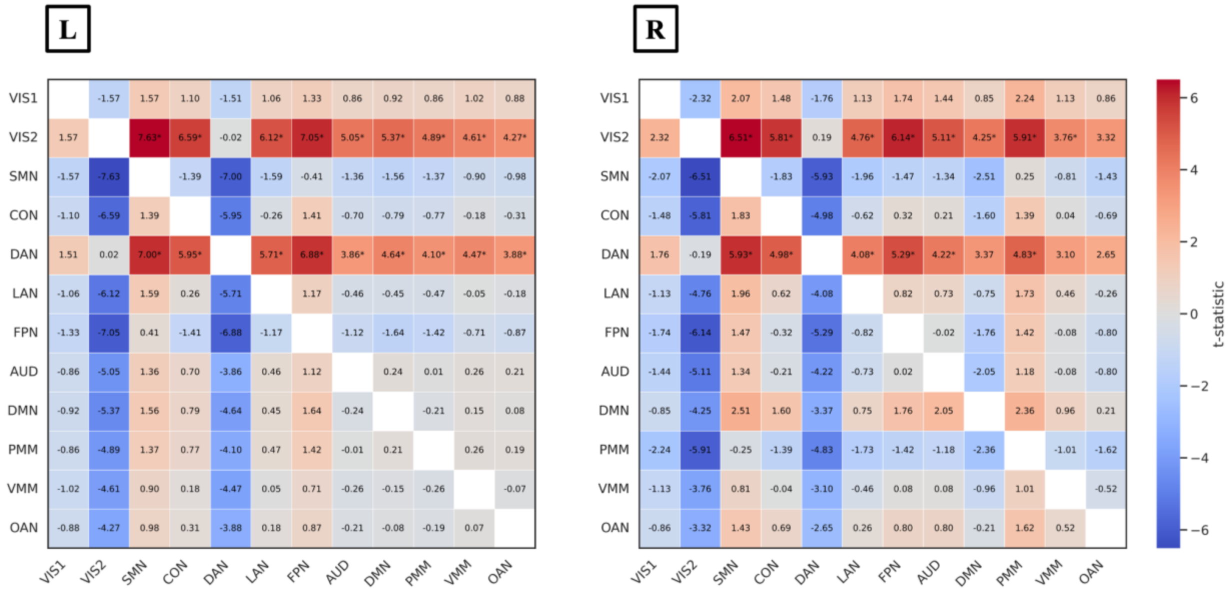

**Figure S5. Full matrix of observed t-statistics for pairwise network comparisons for the left and right hemispheres in the repeat resting-state scan analysis, in comparisons of the contributions of large-scale functional networks.** Each cell reports the t-statistic for the comparison between the row network and the column network (Row minus Column). Positive t-values (shown in shades of red) indicate greater activation in the row network relative to the column network. Negative t-values (shown in shades of blue) indicate greater activation in the column network relative to the row network. The diagonal is masked as it represents self-comparisons.

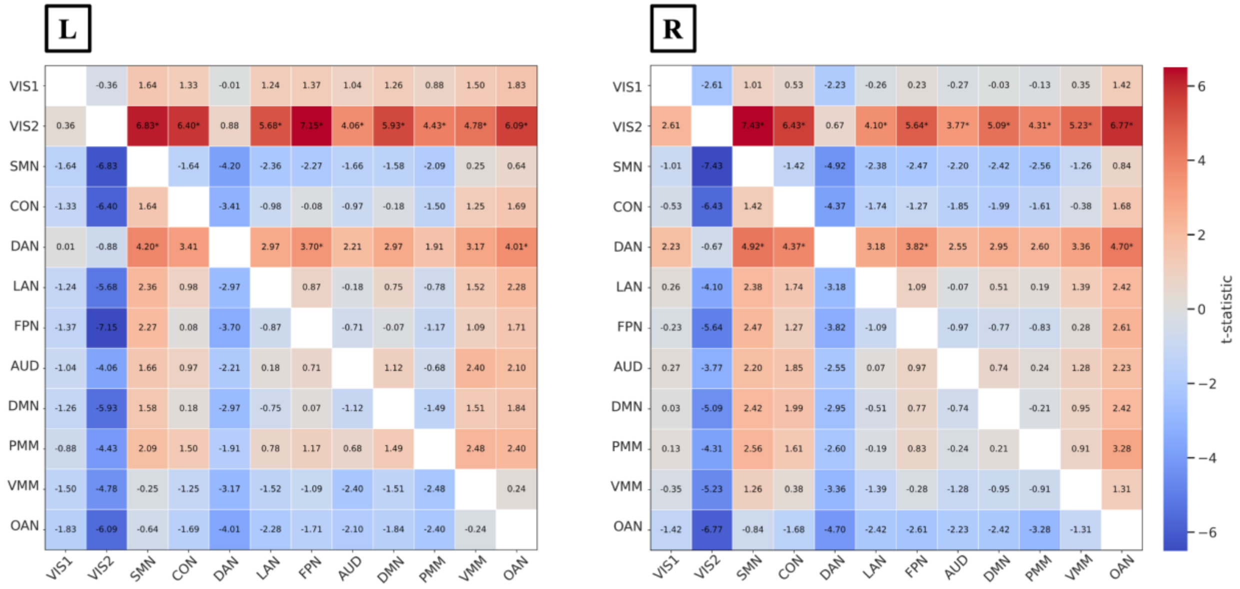

**Table S1.** Full report of the dominance analysis. Estimated partial $R^2$ and Percentages of relative importance in each network (averaged across participants), for CCC in the left and right hemispheres across conditions (response profile).

| | **Left CCC** | | **Right CCC** | | **Left CCC Repeat test** | | **Right CCC Repeat test** | |
|---|---|---|---|---|---|---|---|---|
| **Networks** | Partial $R^2$ | Relative importance | Partial $R^2$ | Relative importance | Partial $R^2$ | Relative importance | Partial $R^2$ | Relative importance |
| **VIS1** | 0.1 | 12.22% | 0.1 | 11.21% | 0.1214 | 14.65% | 0.0941 | 11.15% |
| **VIS2** | 0.21 | 27.00% | 0.22 | 26.72% | 0.2078 | 26.05% | 0.2721 | 32.47% |
| **SMN** | 0.07 | 8.88% | 0.06 | 7.51% | 0.0742 | 9.01% | 0.0572 | 6.55% |
| **CON** | 0.06 | 7.32% | 0.05 | 6.59% | 0.0479 | 5.92% | 0.0519 | 6.19% |
| **DAN** | 0.07 | 9.40% | 0.09 | 10.73% | 0.076 | 9.74% | 0.0776 | 9.25% |
| **LAN** | 0.04 | 5.47% | 0.05 | 6.22% | 0.0424 | 5.59% | 0.0397 | 4.67% |
| **FPN** | 0.05 | 6.61% | 0.06 | 6.70% | 0.0542 | 6.69% | 0.0621 | 7.50% |
| **AUD** | 0.04 | 4.82% | 0.04 | 4.44% | 0.0295 | 3.75% | 0.0497 | 5.89% |
| **DMN** | 0.04 | 5.40% | 0.04 | 4.80% | 0.0414 | 5.33% | 0.0507 | 6.00% |
| **PMM** | 0.04 | 5.01% | 0.05 | 6.66% | 0.0356 | 4.43% | 0.0333 | 4.03% |
| **VMM** | 0.03 | 3.84% | 0.03 | 3.86% | 0.0357 | 4.40% | 0.023 | 2.77% |
| **OAN** | 0.03 | 4.03% | 0.04 | 4.57% | 0.0365 | 4.44% | 0.0303 | 3.52% |
| **Total** | 0.78 | 100.00% | 0.82 | 100.00% | 0.8 | 100.00% | 0.84 | 100.00% |
| **t vs. 0.5** | $t(47) = 17.39$ | | $t(47) = 21.89$ | | $t(26) = 13.11$ | | $t(26) = 20.56$ | |
| **p vs. 0.5** | $4.13 \times 10^{-22}$ | | $2.7 \times 10^{-26}$ | | $5.75 \times 10^{-13}$ | | $1.32 \times 10^{-17}$ | |
